# Implications of critical nodes-dependent unidirectional cross-talk between Plasmodium and Human SUMO pathway proteins in Plasmodium infection

**DOI:** 10.1101/2021.08.18.456755

**Authors:** Ram Kumar Mishra, Jai Shankar Singh, Sajeev T K, Rajlaxmi Panigrahi, Pearl Cherry, Nimisha Abhay Panchakshari, Vaibhav Kumar Shukla, Ashutosh kumar

## Abstract

The endoparasitic pathogen, *Plasmodium falciparum* (Pf), modulates protein-protein interactions to employ post-translational modifications like SUMOylation in order to establish successful infections. The interaction between E1 and E2 (Ubc9) enzymes governs species specificity in the Plasmodium SUMOylation pathway. Here, we demonstrate that a unidirectional cross-species interaction exists between *Pf-*SUMO and Human-E2, whereas Hs-SUMO1 failed to interact with Pf-E2. Biochemical and biophysical analysis revealed that surface-accessible Aspartates of Pf-SUMO determine the efficacy and specificity of SUMO-Ubc9 interactions. Furthermore, we demonstrate that critical residues of the Pf-Ubc9 N-terminal are responsible for the lack of interaction between Hs-SUMO1 and Pf-Ubc9. Mutating these residues to corresponding Hs-Ubc9 residues restore electrostatic, π-π, and hydrophobic interactions and allows efficient cross-species interactions. We suggest that the critical changes acquired on the surfaces of *Plasmodium* SUMO and Ubc9 proteins as nodes can help *Plasmodium* exploit the host SUMOylation machinery. Thus, Pf-SUMO interactions can be targeted for developing antimalarials.

## Introduction

Protein-protein interaction interfaces are crucial for delineating molecular recognition principles, protein association mechanisms, and post-translational modifications (PTM)^1–4^. Parasites often challenge fidelity in target recognition and modifications to develop an infection, survive in the adverse host cellular milieu, and evade the host immune response^5–8^. *Plasmodium falciparum*, known to inflict cerebral malaria in humans, adopts several strategies, including PTMs for infection and survival^9–, 12^. SUMOylation, a post-translational modification similar to ubiquitination, is carried out by a cascade of enzymes, heterodimeric E1 activating (Uba2/Aos1) and E2 conjugating (Ubc9) enzymes, to covalently link the ubiquitin-like polypeptide SUMO to a lysine residue present on the substrate protein. SUMO modified proteins are efficiently deconjugated by sentrin proteases (SENP). SUMOylation pathway enzymes interact non-covalently as well as form a covalent bond with SUMO^13–17^. Detailed studies suggest that non-covalent interactions exist between SUMO and E2 enzymes involving key residues present at the interaction interface^18, 19^. Structural and biochemical studies on the *Plasmodium* SUMOylation pathway suggest that the cognate E1 and E2 enzyme interactions strictly govern the species specificity^11^. Crucial surface residues on E1 and E2 enzymes were mapped as critical nodes governing the pathway-specificity.

Interestingly, disrupting E1-E2 interaction is suggested as an attractive strategy for developing parasite-specific inhibitors^20, 21^. Moreover, studies from various parasites indicate that parasite toxin targets host Ubc9, compromising overall SUMOylation levels for successful infection^22, 23^. *Plasmodium* has elaborate SUMOylation machinery components, and interestingly Pf-SUMO has been found in the cytosol of RBC^9^. *In vitro* results argue that Pf-SUMO can modify a model substrate in the presence of Human E1 and E2 enzymes^10, 11, 24^. Accordingly, small-molecule Pf-SENP1 inhibitor VE-260 affected SUMO processing, inhibited Plasmodium replication, and blocked RBC rupture^25^. These observations motivated our investigation of SUMO-E2 interactions and their relevance in host-pathogen (human-*Plasmodium*) interactions.

Here, we provide a detailed account of Pf-SUMO and Ubc9 interactions and map critical amino acids present on both the proteins as nodes involved in cross-species and species-specific SUMO-Ubc9 interaction. We determined the solution structure of *Plasmodium* SUMO using NMR spectroscopy and suggest that Pf-SUMO and human SUMOs have similar structural arrangements. Next, we delineate a molecular-level understanding of the distinction in interactions between SUMO and E2 proteins by combining biophysical and biochemical approaches. We demonstrate that Pf-SUMO interacts with Hs-E2 albeit with lower affinity; however, Hs-SUMO1 failed to interact with Pf-E2. We also demonstrate that negatively charged Aspartate residues (D68 and D90) of Pf-SUMO, part of the interaction interface, serve as critical nodes and play an essential role in the Pf-SUMO-Ubc9 interaction. Pf-Ubc9 discriminates Pf-SUMO from Hs-SUMO via π-π, hydrophobic and charge-based interactions, mediated by Alanine 13, Glutamate 14, and Alanine 21 residues in the N-terminus of Pf-Ubc9. Combining structural, biochemical, and *in cellulo* observations, we suggest that Pf-SUMO interacts with and utilizes human SUMOylation pathway enzymes. We propose that Pf-SUMO interactions in RBC cytosol can help Plasmodium exploit the host SUMOylation pathway for sustained infection. In addition, targeting Pf-SUMO interactions can help in developing antimalarials.

## RESULTS

### Pf-SUMO displays competitive inter-species interaction with Hs-Ubc9

To develop a precise understanding of host-pathogen (human-*Plasmodium*) interaction, we examined SUMO-Ubc9 interfaces. Alignment of Pf-SUMO against human SUMO paralogs exhibited sequence identities of 41 % with Hs-SUMO1, 45% with Hs-SUMO2, and 46% with Hs-SUMO3 (Supplementary Fig. 1 a). Next, isothermal titration calorimetry (ITC) dependent probing of SUMO-Ubc9 interactions suggested that the binding between Pf-SUMO and Pf-Ubc9 is exothermic and enthalpy-driven, with entropy contributing favorably. The heat exchange profiles fitted well to one-site binding models, indicating single-site binding between the two proteins. The K_d_ value for the Pf-SUMO:Pf-E2 interaction was determined to be 4.05 ± 0.17µM indicating a strong interaction. The K_d_ value of 27.42 ± 1.97µM for Pf-SUMO:Hs-Ubc9 indicated an order of magnitude weaker binding affinity (Fig. 1 a, b and Supplementary Table 1). Similarly, the K_d_ value of 4.18 ± 0.2µM for the Hs-SUMO1:Hs-E2 interaction suggested a robust exothermic mode of binding between them (Supplementary Table 1). Interestingly, no measurable thermodynamic parameters were observed for the Hs-SUMO1:Pf-Ubc9 interaction pair (Fig. 1 c,d). Using Surface Plasmon Resonance, we next compared the binding affinities between Plasmodium and human SUMO proteins with E2 proteins (Hs-Ubc9 and Pf-Ubc9). The binding constant derived herein corroborated with the ITC data, with slight changes in the K_d_ values (Supplementary Fig. 2). The thermodynamic parameters for the interaction between proteins have been summarized in the Supplementary Table 1. Collectively, these findings suggested that Pf-SUMO exhibits intra- as well as inter-species interaction with E2s. However, Hs-SUMO1 exhibited only intra-species interaction with Hs-Ubc9.

**Fig. 1:**
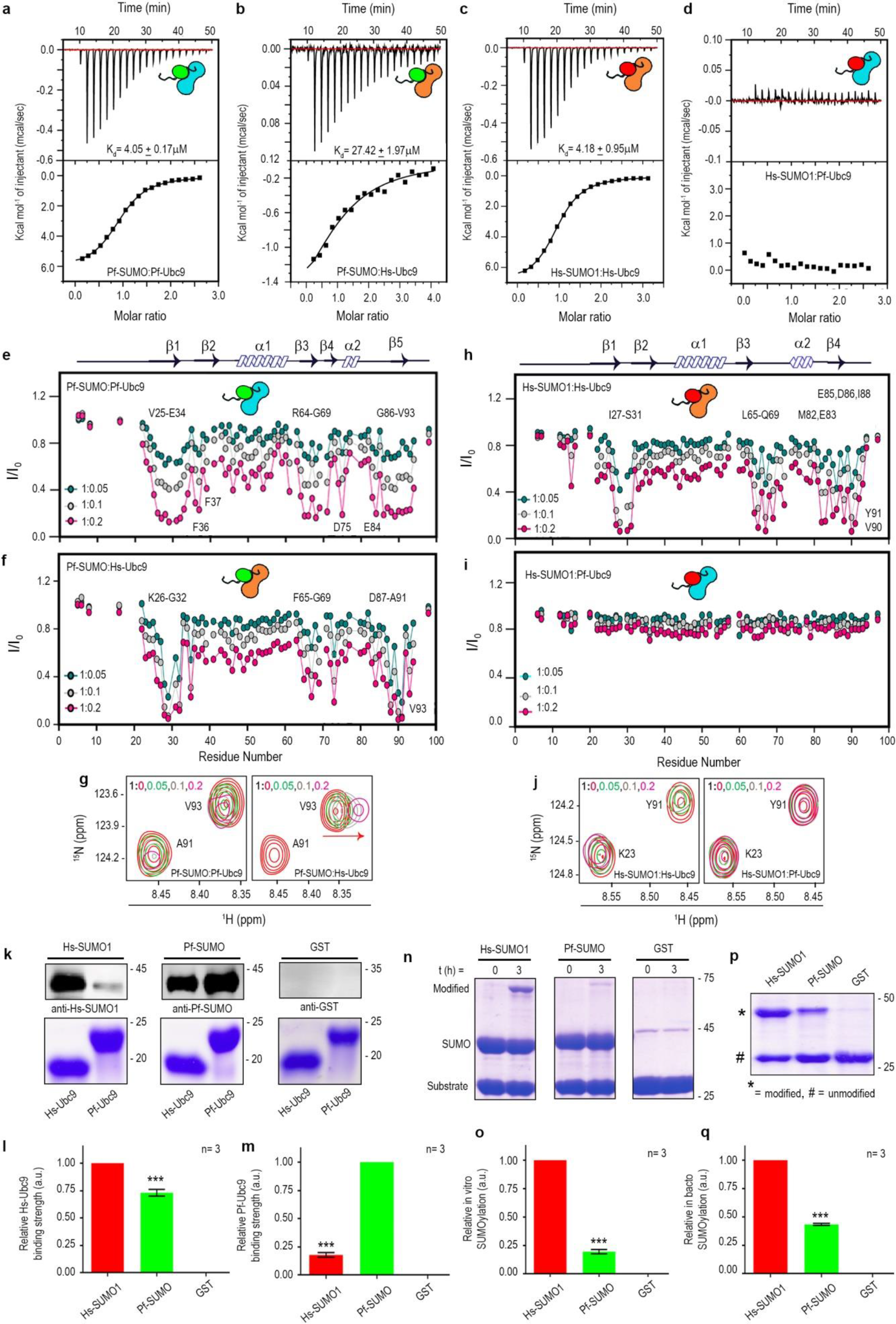
*Plasmodium* SUMO exhibits strong cross-reactivity with human Ubc9 enzyme. Thermodynamic binding analysis utilizing ITC for Pf-SUMO with Pf-Ubc9 **(a)**, Pf-SUMO with Hs-Ubc9 **(b)**, Hs-SUMO1 with Hs-Ubc9 **(c)**, and Hs-SUMO1 with Pf-Ubc9 **(d)**. The upper panels represent the raw data, and the corresponding lower panels indicate the curve fitting of the graph to extract thermodynamic parameters. Intensity profile (I/I_0_) of amide cross-peaks obtained from ^15^N-^1^H HSQC spectra at different ratios of SUMO and Ubc9 (1:0.05, dark cyan; 1:0.1, gray; 1:0.2 pink) for Pf-SUMO-Pf-Ubc9 **(e)** and Pf-SUMO-Hs-Ubc9 **(f)** pairs. **(g)** Excerpts of selected peaks from Pf-SUMO interactions as observed in (e) and (f). Similar analysis as in **e** and **f** but Hs-SUMO1 with Hs-Ubc9 **(h)** and Hs-SUMO1 with Pf-Ubc9 **(i)**. **(j)** Excerpts of selected peaks from Hs-SUMO1 interactions as observed in (h) and (i). Overlay ^15^N-^1^H heteronuclear single-quantum coherence (HSQC) spectra of Pf-SUMO with Ubc9 enzymes at a ratio of 1: 0.0, 0.05,, 0.2, i.e. free state (black colour) to the bound state. Arrow indicates the direction of chemical shift perturbations. **(k)** *In vitro* binding assay for Pf-SUMO and Hs-SUMO1 with human and plasmodium Ubc9s. GST is used as a negative control for binding analysis. Upper panels show western blotting with respective α-SUMO antibodies, and lower panels represent coomassie stained gels showing Ubc9s used for pulldown. The quantification of SUMO binding affinities as seen in (k) with Hs-Ubc9 **(l)** and Pf-Ubc9 **(m)**. **(n)** *In vitro* SUMOylation with Hs-SUMO1 and Pf-SUMO in the presence of purified human SUMOylation machinery components and a standard peptide substrate and the quantification of the same in (n) **(o)**. GST serves as a negative control. *In bacto* SUMOylation with Hs-SUMO1 and Pf-SUMO in the presence of human SUMOylation machinery and a standard peptide substrate expressed inside bacteria **(p)** and the quantification of the same in (p) **(q)**. GST serves as a negative control. All statistical analysis was carried out using GraphPad Prism 8.4.3. Column analysis of data sets carried out by One-way ANOVA (nonparametric). Dunnett’s test was used for multiple comparisons. Family-wise significance and confidence level is p<0.001. Simple cartoons above relevant panels represent Hs-SUMO1 (red), Pf-SUMO (green), Hs-Ubc9 (orange), and Pf-Ubc9 (cyan).

We next explored the residue-specific information on the interactions between Pf-SUMO and Hs-SUMO1 with the Ubc9 enzymes. We purified these proteins (Supplementary Fig. 1 b) and recorded a set of 2D ^15^N-^1^H heteronuclear single quantum coherence (HSQC) spectra on ^15^N labeled Pf-SUMO (250 µM) and then titrated with increasing concentration (0.05, 0.1, 0.2, 0.4, 0.6 molar equivalent) of unlabeled Pf-Ubc9 and Hs-Ubc9. Position of resonance peaks may shift or reduce in intensity due to protein-protein interaction. Further, chemical shift perturbation (CSP) and effects of protein-complex tumbling indicate binding strengths^26, 27^. For Pf-SUMO’s titration with Pf-Ubc9 (Supplementary Fig. 3 a), most cross-peaks disappeared, and very few cross-peaks shifted when Pf-Ubc9 concentration was increased from 0.05 to 0.2 molar ratio. A significant decrease in peak intensities was observed for the residues V25-A33 and F37 located at the structured region of N-terminus, R64-G69, R71, and D75 in the middle region G86-V93 in the C-terminus of Pf-SUMO (Fig. 1 e and Supplementary Fig. 3 a). Beyond a molar ratio of 0.2, most peaks disappeared, indicating a stronger binding between proteins in the intermediate exchange regime at the NMR time-scale^27^. Similarly, for Pf-SUMO’s titration with Hs-Ubc9, we observed a considerable shift or disappearance in resonances of a significant number of cross-peaks. The prominent regions with intensity decrease and/or CSP were clustered around Pf-SUMO residues K26-G32, V35 in the N-terminus, F65-G69, H73 in the middle, and D87-A91 in the C-terminus (Fig. 1 f and Supplementary Fig. 3 b). Interestingly, the same region of Pf-SUMO appeared to show interaction with the Hs-Ubc9 barring few residues, which may be critical for differential interaction strength. The excerpts of the intra- and interspecies interaction of Pf-SUMO are shown in Fig. 1 g. Pf-SUMO:Hs-Ubc9 interactions demonstrated significant CSPs suggesting that these interactions occur in an intermediate to fast exchange regime at the NMR time scale^28^. Similarly, the titration experiments for Hs-SUMO1 with Ubc9 enzymes suggested strong interaction with most of the cross-peaks disappearing and observation of CSPs for very few cross-peaks at different titration ratios (Fig. 1 h and Supplementary Fig. 3 c). A significant decrease in the intensity due to line broadening was observed in similar regions (viz, N-terminus, middle and C-terminal region) of Hs-SUMO’s, as seen in Pf-SUMO:Pf-Ubc9 interaction. Interestingly, for Hs-SUMO1:Pf-Ubc9 interaction, neither a peak shift nor peak disappearance was observed (Fig. 1 i and Supplementary Fig. 3 d). The excerpts from the intra- and interspecies interaction of Hs-SUMO1 are shown in Fig. 1 j.

Next, we asked if these biophysical assessments of intra- and interspecies SUMO-Ubc9 interactions were biochemically and functionally relevant. We immobilized SUMO on an affinity column and passed E2 proteins over the beads to assess *in vitro* binding. Protein pulldown assays demonstrate strong intra-species interaction between SUMO and Ubc9. In assays of inter-species SUMO-Ubc9 interactions, the Pf-SUMO was observed to bind Hs-Ubc9 with significant affinity. In contrast, Hs-SUMO1 failed to display appreciable interactions with Pf-Ubc9 (Fig. 1 k). Quantification of these observations suggests that the relative affinity of the Pf-SUMO:Hs-Ubc9 interaction is ∼67%, in contrast to the ∼12% relative binding affinity of Hs-SUMO1:Pf-Ubc9 (Fig. 1 l,m). Next, we assessed the functionality of Pf-SUMO:Hs-Ubc9 interaction through an *in vitro* and *in bacto* (SUMOylation inside bacteria) SUMOylation reaction in the presence of SUMO machinery and a standard substrate (Supplementary Fig. 4). The *in vitro* and *in bacto* SUMOylation reactions suggested that using human SUMOylation machinery, Pf-SUMO can SUMOylate the substrate but at a ∼five-fold lesser efficiency than Hs-SUMO1 (Fig. 1 n-q). In comparison, Pf-SUMO mediated *in bacto* SUMOylation was ∼two-fold more efficient than *in vitro* SUMOylation, probably due to the prolonged duration of the SUMOylation reaction. Together, the biophysical (NMR, ITC, and SPR), biochemical (pulldown), and functional (SUMOylation) analyses establish significant interaction between Pf-SUMO:Hs-Ubc9.

### Pf-SUMO shares structural conservation with Hs-SUMO paralogs and Ubc9 interaction interface

Several protein-protein complexes are characterized by shape complementarity recognition at the interface (Supplementary Fig. 5 a). Knowing Pf-SUMO’s three-dimensional structure is necessary to precisely understand the details of interactions that exist during cross-species recognition of Pf-SUMO and Hs-SUMO1 by Ubc9. Using conventional NMR experiments (as detailed in the method section) and distance restraints, we determined the 3D structure of Pf-SUMO in solution. The cartoon representation of the lowest energy structure (Supplementary Fig. 6 a,b) and the wire representations of the final ensemble of Pf-SUMO’s ten structures are shown in (Fig. 2 a). The pairwise RMSD for the ordered regions (aa 22-98) of the final ensemble of the ten lowest energy structures was 0.61 Å. The atomic coordinates for all ten Pf-SUMO protein structures have been deposited in the PDB (PDB code: 5GJL). The rest of the structural parameters and quality of the Pf-SUMO structure is summarized in (Supplementary Table 2). Briefly, the solution structure of Pf-SUMO has a conserved SUMO fold, consisting of a four-stranded mixed β-sheet, one helix, and one helical turn. The strand ordering is β2-β1-β4-β3, in which the two central strands (β1-β4) are parallel and strands (β2-β1 and β3-β4) run antiparallel to each other (Fig. 2 b).

**Fig. 2:**
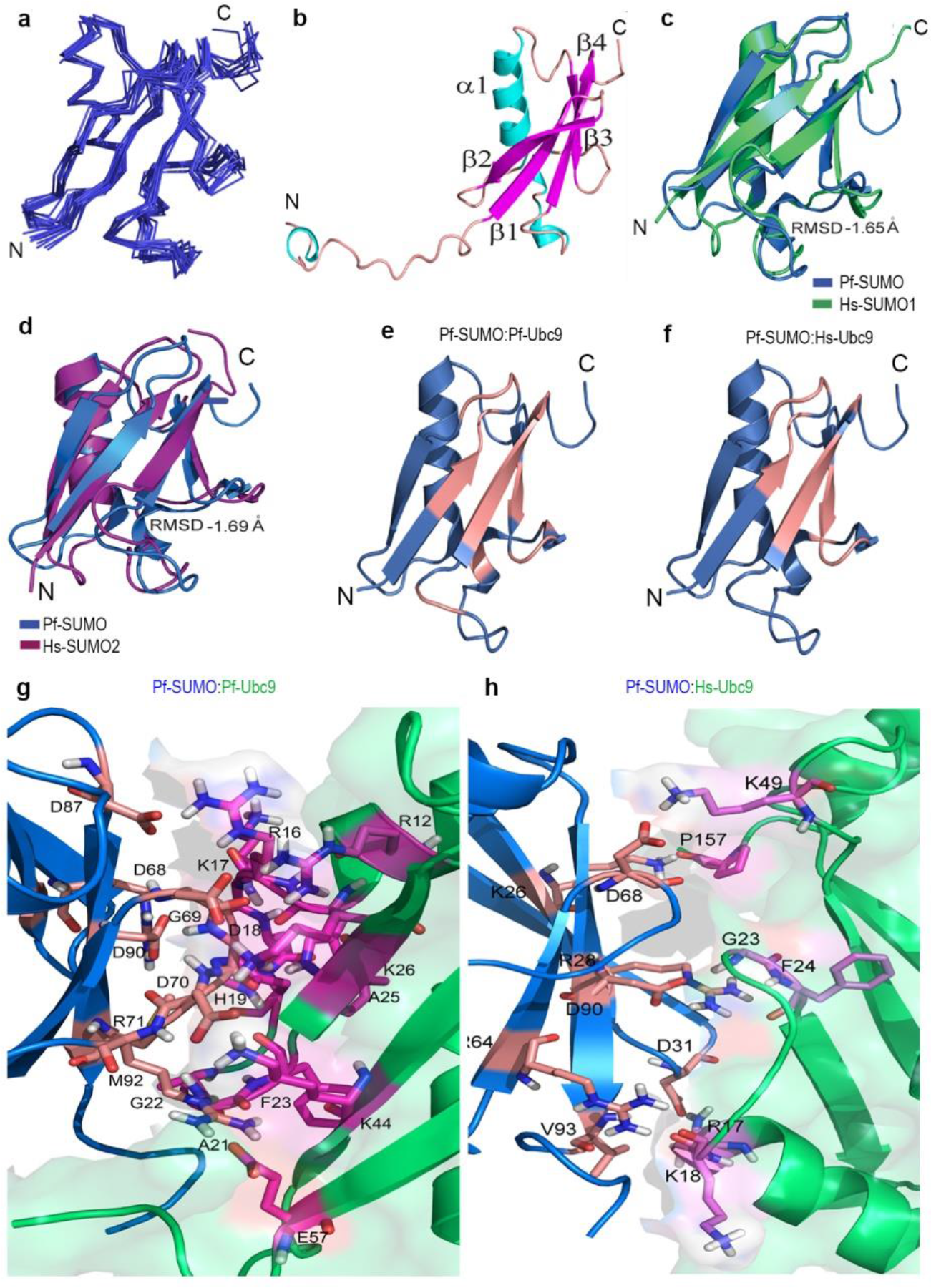
NMR-derived solution structure of Pf-SUMO and molecular docking of Pf-SUMO protein with E2 enzymes. **a)** Superimposition of backbone traces from the final ensemble of 10 structures with lowest target function from 22-98 residues. **b)** Cartoon diagram representing the lowest energy structure of Pf-SUMO. The individual β-strands and α-helices are labeled. The β-strands are β1 (I23-V27), β2 (V35-I39), β3 (V63-L66), and β4 (D87-V93); and α-helices are α1 (L45-L56). **c, d)** Overlap of average Pf-SUMO structure over Hs-SUMO1 and Hs-SUMO2 respectively and calculated RMSD mentioned aligned overlapped structures. **e, f)** Surface conservation of Pf-SUMO while binding with E2 enzymes. Salmon colour in Pf-SUMO structure represents the surface residues interacting with Pf-Ubc9 and Hs-Ubc9 enzymes, respectively, as determined using NMR. **g, h)** Docked model of Pf-SUMO residues showing interaction with the residues of Pf-Ubc9 and Hs-Ubc9 enzymes, respectively. The colour coding for Pf-SUMO is blue colour whereas Pf-Ubc9 and Hs-Ubc9 are green colour respectively. Interacting residues of Pf-SUMO are in salmon colour, and Pf-Ubc9 and Hs-Ubc9 are in magenta, respectively.

We evaluated the structural differences between Pf-SUMO and Hs-SUMO paralog structures and superimposed the NMR structure of Pf-SUMO over those of human SUMO1 (PDB ID: 2N1V, RMSD = 1.65 Å) and human SUMO2 (PDB ID: 2N1W, RMSD = 1.69 Å), respectively (Fig. 2 c,d). The Pf-SUMO showed high structural alignment with the Hs-SUMO1. Even though the overall fold was the same for these proteins, small structural differences between the structure of Pf-SUMO and Hs-SUMO1/2 appeared in the loop regions: R28-D31 (between β1-β2), D68-D70 (between β3-β4), D75 (α2 helical turn), and D85-D87 (in the β5 region of C-terminus).

Detailed assessment of SUMO-Ubc9 complexes showed that the interaction between these proteins is primarily charge-dependent, where the positively charged N-terminal of Ubc9 interacts with the negatively charged C-terminal pocket of SUMO^19, 29^ (Supplementary Fig. 5 b). In the previous section, we have reported distinct regions of interaction between SUMO and Ubc9. All Pf-SUMO residues involved in interaction with Pf-Ubc9 and Hs-Ubc9 enzymes have been demarcated in salmon colour (Fig. 2 e,f). The NMR-derived structure of Pf-SUMO was docked with Ubc9 proteins using the HADDOCK 2.2 server. The docked complex structures were stabilized by a series of H-bonds as well as salt bridges. The structural details of the interface residues of Pf-SUMO with Ubc9 enzymes have been summarized (Supplementary Fig. 6 c,d). Upon comparing the docked complex of Pf-SUMO:Pf-Ubc9 with that of the Pf-SUMO:Hs-Ubc9, the electrostatic interactions and the H-bond contacts at the interface were found to be 50% higher for the former complex (Fig. 2 g,h and Supplementary Table 3), which explains the reason behind the strong interaction of Pf-SUMO with the Pf-Ubc9 compared to Hs-Ubc9. Eventually, molecular docking substantiated NMR observation that the negatively charged residues at positions D68, G69, D87, and D90 in Pf*-*SUMO, forming a stable complex with positively charged residues of Ubc9.

### C-terminal region of Pf-SUMO is critically involved in the interaction with Ubc9

SUMO interacts non-covalently on a site located at the back-side of the Ubc9^19, 29^. NMR titration experiments indicated three prominent patches on Pf-SUMO (shown on primary sequence) required for interaction with Pf-Ubc9 or Hs-Ubc9 (Fig. 3 a). Two of these patches are present in the C-terminal region centered around D68 and D90 residues. To establish the importance of Pf-SUMO regions required for interaction with Ubc9, we generated chimeras between Pf-SUMO and human SUMO1. The Pf-N chimera had amino acid 1-60 of Pf-SUMO, amino acid 61-97 of Hs-SUMO1, and the Pf-C chimera had amino acids 1-61 of Hs-SUMO1 and residues 61-98 of Pf-SUMO (Fig. 3 b). Pulldown of chimera proteins on Hs-Ubc9 and Pf-Ubc9 demonstrated that Pf-N chimera exhibits stronger interactions with Hs-Ubc9 (Fig. 3 c,d) while the Pf-C chimera did the same with Pf-Ubc9 (Fig. 3 e,f), indicating a role for the C-terminus in recognition of the cognate partner. These biochemical results corroborate the findings from the NMR data and identify two out of the three Ubc9 interacting regions in the Pf-SUMO C-terminal half. Further, the *in vitro* SUMOylation reactions with SUMO chimera proteins indicated a significantly improved SUMO modification with Pf-N chimera compared to the Pf-C chimera (Fig. 3 g,h). While strongly supporting the importance of C-terminal residues of SUMOs in cognate Ubc9 recognition and substrate modification, these results also hint at the involvement of N-terminal residues.

**Fig. 3:**
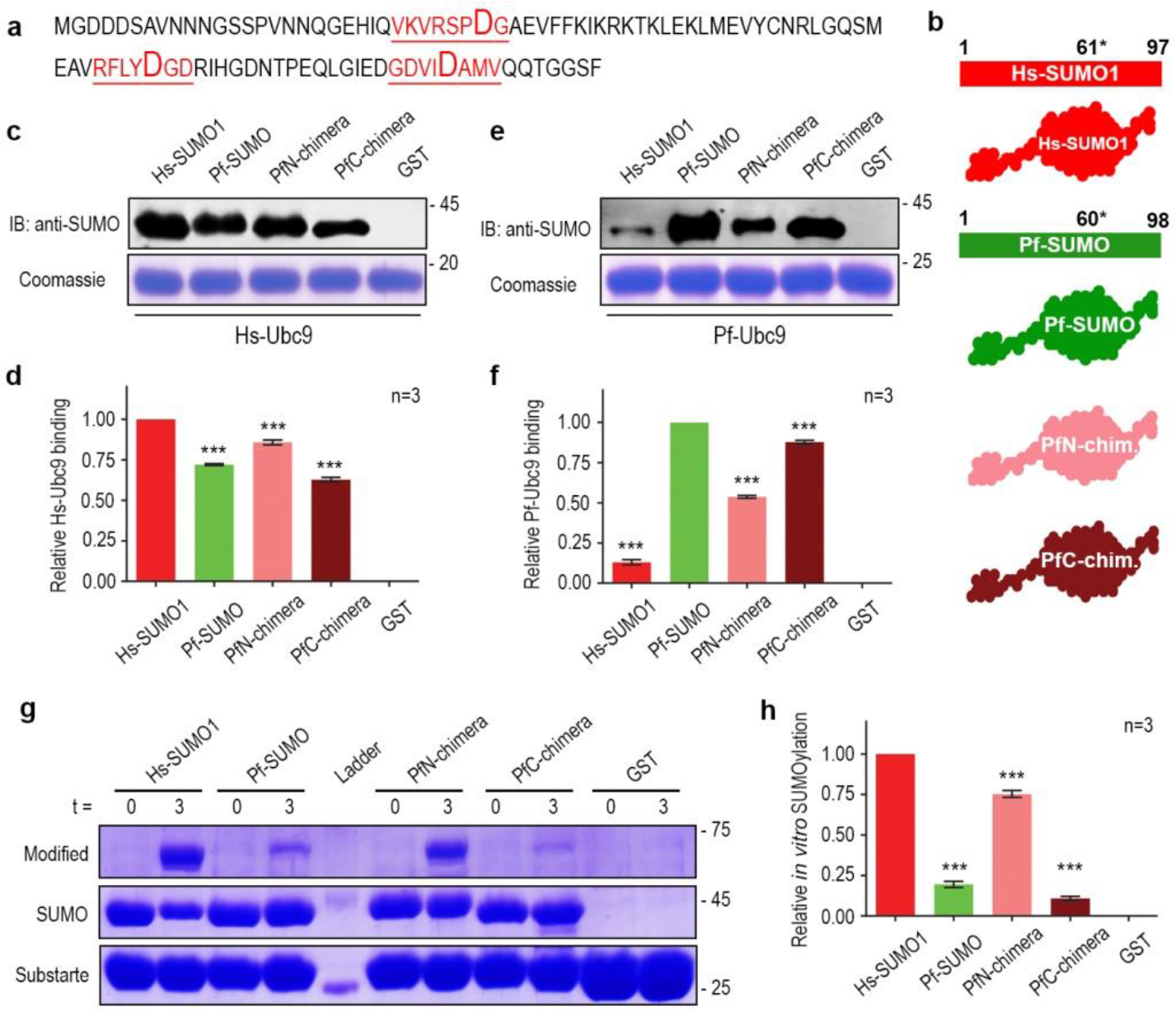
Contributions from both the amino- and carboxy terminus of Pf-SUMO facilitates cross-interaction with Hs-Ubc9. **a)** Primary sequence of *Plasmodium* SUMO highlighting Ubc9 interacting patches as determined from NMR titration analysis **b)** Schematic representation of Chimera generation scheme and cartoon diagram of SUMO chimera proteins. * (61* and 60*) indicates the position at which the protein has been divided into two halves to generate the chimera (PfN, carrying N-term of Pf-SUMO and PfC, carrying C-term of Pf-SUMO) proteins. **c)** *In vitro* binding of chimera proteins over Hs-Ubc9. Upper panels indicate anti-SUMO antibody blotting, and the coomassie stained lower panels indicate Hs-Ubc9 levels. **d)** The graphical representation of quantification of SUMO binding as seen in (c). **e and f)** Same as in (c) and (d), respectively, but the pulldown is performed on Pf-Ubc9. **g)** *In vitro* SUMOylation with chimera proteins in the presence of purified human SUMOylation machinery components and a standard peptide substrate. GST serves as a negative control for the reaction. **h)** The quantification of the observation made in (g). All experiments are performed at least three times independently. All statistical analysis was carried out using GraphPad Prism 8.4.3. Column analysis of data sets carried out by One-way ANOVA (nonparametric). Dunnett’s test was used for multiple comparisons. Family-wise significance and confidence level is p<0.001.

### Conserved Aspartate residues are critical for SUMO-Ubc9 interaction

Several analyses, including the comprehensive mutational analysis of yeast Smt3, report the importance of C-terminal Aspartate residues in Smt3-Ubc9 interaction, where mutations in these negatively charged residues of Smt3 abolished interaction with Ubc9 and induced lethality in yeast^30^. Our NMR titration studies suggested three binding patches centered around K26-D31, Y67-D70, and E84-A91 residues of Pf-SUMO engaged in Ubc9 interaction (Fig. 1 e,f), where residue D68 and D90 were first to show intensity decay during titration. Moreover, molecular docking studies analyzing the interaction of Ubc9s with Pf-SUMO reestablished criticality of Aspartates at position 68 (D68) and at position 90 (D90) in the carboxy-terminal of the Pf-SUMO to be involved in the stabilization of protein complexes.

Point mutants for D31, D68, and D90 residues were generated to validate the criticality of these residues. These residues were mutated to Alanine; and, D68 and D90 were mutated to Lysine also. We probed the aspartate (D31, D68, and D90) mutants of Pf-SUMO for their importance in regulating the interaction with Ubc9s. The quantification of data from *in vitro* binding with Ubc9s, *in vitro,* and *in bacto* SUMOylation analysis for Pf-SUMO D31A mutant did not show any observable differences compared with the Pf-SUMO (Supplementary Fig. 7 a-h). Therefore, we focused on analyzing the aspartate residues lying in the C-terminal half of the Pf-SUMO. We performed *in vitro* pulldown assays with Pf-SUMO D68A, D68K, D90A, and D90K mutants and observed that in comparison to Pf-SUMO, the aspartate mutants, Pf-SUMO D68A, D68K, D90A, and D90K exhibited a significant reduction (∼90%) in their ability to interact with Hs-Ubc9, with D90 mutants maximally impairing (Fig. 4 a,b). Similarly, in pulldown assay with Pf-Ubc9, the D68A, D68K, and D90K mutants of Pf-SUMO showed a significant reduction (80-90%) than Pf-SUMO. Intriguingly, the D90A mutant was unperturbed in its ability to bind with Pf-Ubc9 (Fig. 4 c,d). A single site binding with the K_d_ value of 2.50 ± 0.97 µM was observed for the Pf-SUMO D90A:Pf-Ubc9 interaction, indicating an unperturbed interaction matching Pf-SUMO:Pf-Ubc9 levels (Fig. 4 e). SPR thermodynamic parameters showed a similar binding constant (K_d_ 3.02 ± 0.93 µM) between these systems (Supplementary Fig. 8 g). The dissociation constant between D90A and Hs-Ubc9 was found to be ∼four-fold weaker than the wildtype (data not shown).

**Fig. 4:**
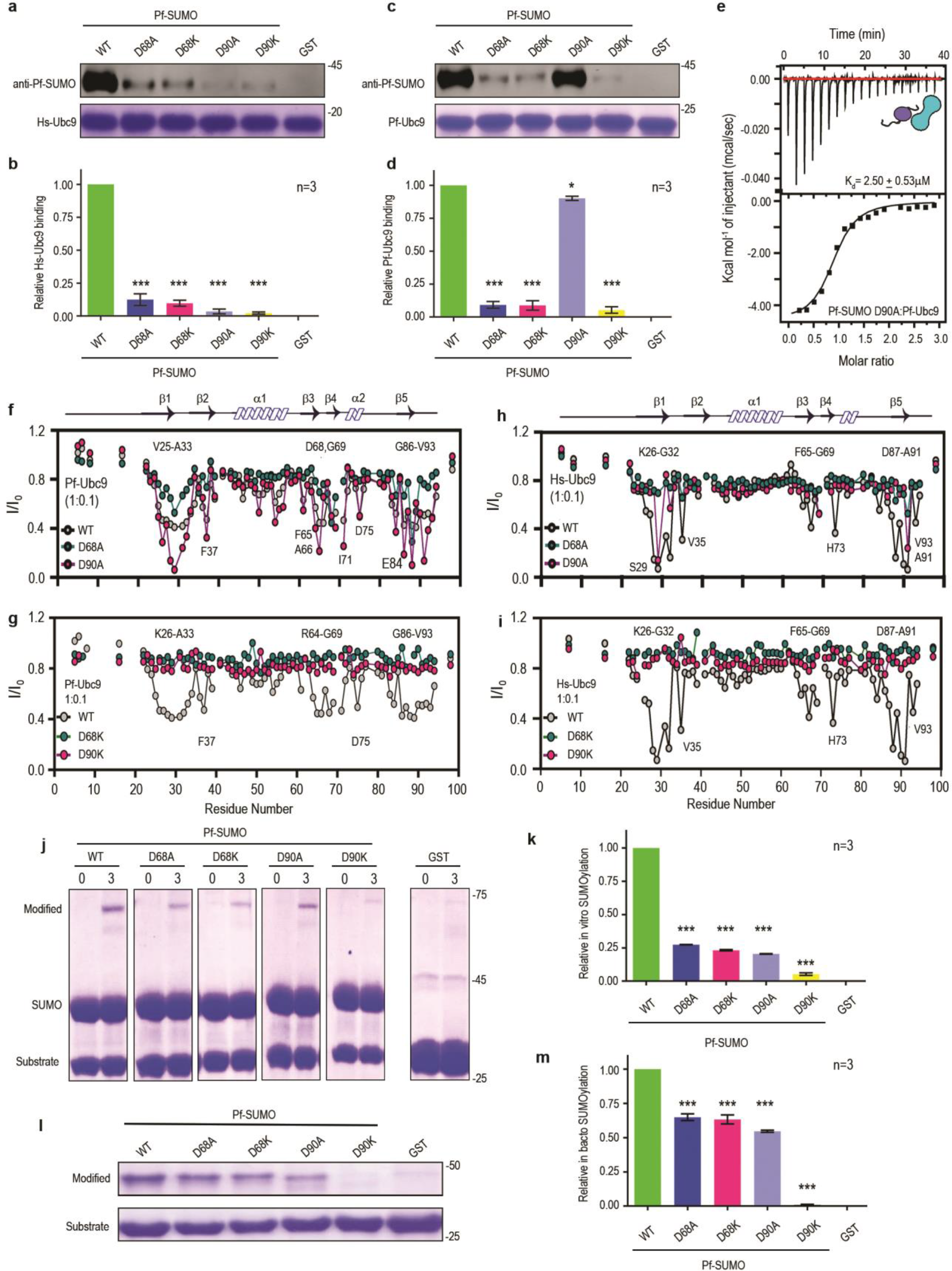
Negatively charged nodes at Aspartate 68 and 90th position govern specific Pf-SUMO interactions with Ubc9. *In vitro* binding assay of Pf-SUMO mutant proteins over Hs-Ubc9 **a)**. Upper panels indicate anti-Pf-SUMO antibody blotting, and the coomassie stained lower panels indicate Hs-Ubc9 levels. **b)** The quantification of Pf-SUMO binding as seen in (a). **c and d)** Same as in a and b; however, the binding experiments are performed over the Pf-Ubc9. GST serves as a negative control. **e)** Thermodynamic binding analysis using ITC for Pf-SUMO D90A mutant with Pf-Ubc9. The upper panels represent the raw data, and the corresponding lower panels indicate the curve fitting of the graph. **f)** Intensity profile (I/I_0_) of amide cross-peaks obtained from ^15^N-^1^H HSQC spectra at 1:0.1 ratios of Pf-SUMO wild type and indicated mutants (Gray; wild type, Green; Pf-SUMO-D68A, Pink; Pf-SUMO-D90A) with Pf-Ubc9. Arrow indicates the direction of chemical shift perturbations. **g)** Same as in f, but D68K and D90K mutants of Pf-SUMO along with wildtype were used in the interaction with Pf-Ubc9. **h and i)** Same as in (f and g); however, Hs-Ubc9 was used for interaction analysis. **j)** *In vitro* SUMOylation with Pf-SUMO wildtype and indicated mutants in the presence of purified human SUMOylation machinery components and a standard peptide substrate. GST serves as a negative control. **k)** The quantification of the binding as observed in (j). **l)** *In bacto* SUMOylation with Pf-SUMO wildtype and indicated mutants in the presence of human SUMOylation machinery components and a standard peptide substrate expressed inside bacteria. GST serves as a negative control. **m)** The quantification of the binding as observed in (l). All experiments are performed at least three times independently. All statistical analysis was carried out using GraphPad Prism 8.4.3. Column analysis of data sets carried out by One-way ANOVA (nonparametric). Dunnett’s test was used for multiple comparisons. Family-wise significance and confidence level is p<0.001. The cartoon inside figure 4(e) represents Pf-SUMO mutants and Pf-Ubc9 (cyan).

We recorded a series of 2D {^15^N-^1^H} HSQC spectra on the ^15^N labeled Pf-SUMO (WT) and its mutants (D68A/K and D90A/K) in the free form and the complex state formed during titration with Pf-Ubc9 or Hs-Ubc9. Next, we probed the effect of Pf-SUMO charge neutralization mutants at the 68^th^ and 90^th^ positions on Ubc9 binding. At the 0.1 molar ratio of Pf*-*Ubc9, significant CSPs for the D68A were observed, confirming a weaker binding than the wildtype protein (Supplementary Fig 8 a-c). For the D90A mutant, at a similar molar ratio of 0.1 Pf-Ubc9, substantial line broadening leading to a significant decrease in intensity was observed in the established three prominent regions of Pf-SUMO. Furthermore, the decrease in peak intensities for the D90A mutant was ∼1.5-fold higher than that of the wildtype, confirming an equal or even stronger binding than the wild type. However, the titration analysis for D90A and D68A mutants with Hs-Ubc9 did not show much CSPs other than S29 and A91 residues, suggesting a much weaker or no binding (Fig. 4 f,h, Supplementary 8 e). Next, charge reversal mutants D68K and D90K also neither show CSPs nor intensity change upon titration with Pf-Ubc9 or Hs-Ubc9, indicating critical positioning of these negatively charged residues required for interaction with Ubc9s (Fig. 4 g,I, Supplementary 8 d,f).

Next, we examined the effect of compromised interaction of the D68 and D90 mutants to SUMOylate using the host SUMOylation machinery. *In vitro* SUMOylation with D68A, D68K, D90A, and D90K mutants showed a significant reduction in their ability to modify the substrate (Fig. 4 j,k). Expectedly, the *in bacto* SUMOylation outcomes also indicated a significant reduction in their ability to modify the substrate (Fig. 4 i,m). The Pf-SUMO D68A, D68K, and D90A mutants resulted in ∼three-fold impairment in their SUMOylation abilities; the D90K mutant appeared inactive with near-zero SUMOylation levels. The extent of decreased SUMOylation, under *in vitro* and *in bacto* SUMOylation reaction conditions, observed with these Pf-SUMO mutants corroborated very well with the reduction in their ability to interact with Hs-Ubc9. Taken together, we demonstrate that the Aspartates at positions 68^th^ and 90^th^ are critical for Ubc9 interaction.

### The divergent N-terminal region of Pf-Ubc9 governs species-specific interactions with Pf-SUMO

Structural and biochemical analyses have demonstrated that the N-terminal helix of Ubc9 is a major player in the non-covalent interaction with SUMO^18, 19^. Further, the N-terminal region of Pf-Ubc9 (aa 1-81) contributed towards species-specificity of E1-E2 interaction^10^. Moreover, our initial results suggested that the Pf-Ubc9 does not interact with Hs-SUMO1. Therefore, we asked if the sequence diversity in the N-terminal region of the Pf-Ubc9 does not allow Hs-SUMO1 binding. While Pf-Ubc9 and Hs-Ubc9 exhibited an overall 61% identity (Supplementary Fig. 9 a), alignment of the first 25 amino acids from the N-terminus of Ubc9 from different eukaryotic animals indicated that Pf-Ubc9 has diverged at specific residues (Supplementary Fig. 9 b). With Alanine, Glutamate, and Alanine at positions 13, 14, and 21, respectively, a clear divergence can be observed in the SUMO binding pocket residues of Pf-Ubc9. In comparison, Hs-Ubc9 had Lys, Ala, and Phe residues at equivalent positions. Thus, we reasoned that the nature of these amino acids at critical nodes of Ubc9 determines the intra- and interspecies interaction with SUMO. Molecular docking analyses of the SUMO-Ubc9 complex structures clearly demonstrated that electrostatic forces stabilize the interface residues. The Y91 residue of Hs-SUMO1 forms π-π stacking with the F22 residue of Hs-Ubc9 is also supported by neighboring residues electrostatically (Fig. 5 a). Though Pf-SUMO:Pf-Ubc9 was stabilized by electrostatic and hydrophobic interactions only, this crucial π-π stacking was missing. To delineate the importance of the interaction between these residues, we mutated alanine and glutamate the 13^th^ and 14^th^ positions into lysine and alanine, respectively, in the Pf-Ubc9 (Pf-Ubc9 KA mutant) and the 21^st^ position residue into phenylalanine (A21F mutant), and a combination of these three mutations (KAF mutant). With these KAF mutations in the N-terminus, Pf-Ubc9 resembles Hs-Ubc9. The NMR titration experiments for ^15^N Hs-SUMO-1 with Pf-Ubc9 A21F mutant identified the same three binding sites reported in earlier sections (Fig. 5 b,c, Supplementary Fig. 9 e,f). Though the interaction strength was weaker compared to the Pf-SUMO:Pf-Ubc9 interaction, CSPs were shown by many residues. Furthermore, molecular docking identified a stabilized interface between Hs-SUMO1 and Pf-Ubc9 KAF mutant (Fig. 5 d and Supplementary Fig. 9 c,d).

**Fig. 5:**
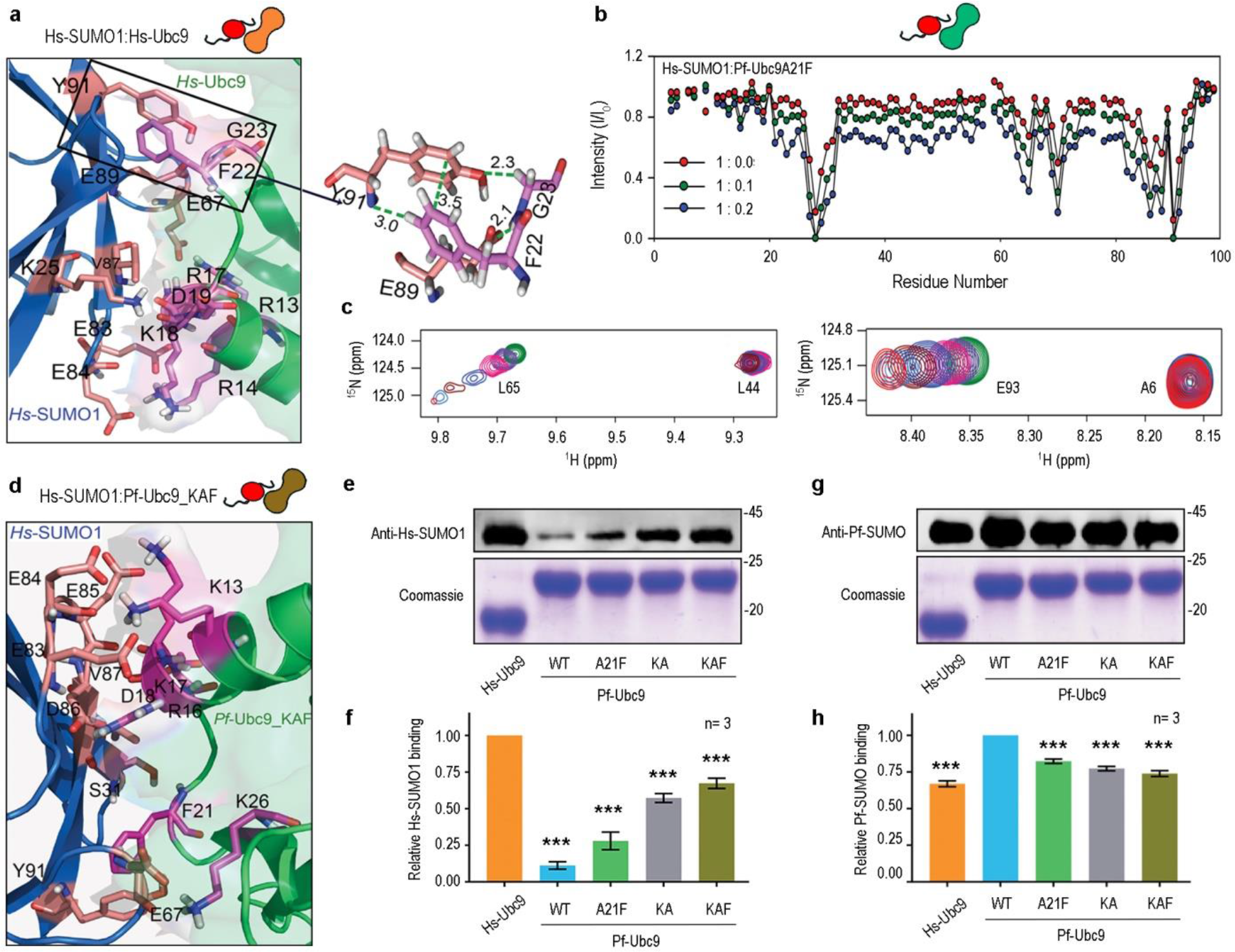
N-terminal hydrophobic residues of Pf-Ubc9 are critical regulators of cross-species interactions. **a)** Docked model of Hs-SUMO1 residues showing interaction with the residues of Hs-Ubc9. The colour coding for Hs-SUMO1 and Hs-Ubc9 are blue and green, respectively. The boxed region is further zoomed in to show the π-π and H-bonding interaction between Y91 of Hs-SUMO1 and F22 of Hs-Ubc9 enzyme. **b)** Overlay and intensity profile (I/I_0_) of amide cross-peaks obtained from ^15^N-^1^H HSQC spectra of Hs-SUMO1 in the presence of different equivalents of A21F Pf-Ubc9, respectively. **c)** Excerpts of selected peaks from Pf-Ubc9A21F interaction with Hs-SUMO1 as observed in (b). **d)** Docked model of Hs-SUMO1 residues showing interaction with the residues of Pf-Ubc9 KAF triple mutant. The colour coding for Hs-SUMO1 and mutant Pf-Ubc9 are respectively blue and green. **e)** *In vitro* binding assay of *Hs*-SUMO1 over Pf-Ubc9 mutants. Upper panels indicate anti-Hs-SUMO1 antibody blotting, and the coomassie stained lower panels indicate Pf-Ubc9 protein levels. Hs-Ubc9 is used as a positive control in the reaction. **f)** The quantification of Hs-SUMO1 binding as seen in (e). **g)** Same as in (e); however, the Pf-SUMO binding is performed over the Pf-Ubc9 mutants. Upper panels indicate anti-Pf-SUMO antibody blotting, and the coomassie stained lower panels indicate Ubc9 levels. **h)** The quantification of Hs-SUMO1 binding as seen in (g). All experiments are performed at least three times independently. All statistical analysis was carried out using GraphPad Prism 8.4.3. Column analysis of data sets carried out by One-way ANOVA (nonparametric). Dunnett’s test was used for multiple comparisons. Family-wise significance and confidence level is p<0.001. Simple cartoons above relevant panels represent Hs-SUMO1 (red) and Pf-Ubc9 mutants.

In parallel, pulldown experiments with these Pf-Ubc9 mutants yielded a significant increase in their interaction with Hs-SUMO1. Compared to the poor interaction observed between Hs-SUMO1:Pf-Ubc9 (∼10% of Hs-SUMO1:Hs-Ubc9 interaction strength), the A21F mutation improved interaction by ∼2.5-fold to register ∼28% interaction strength, and the KA and KAF mutants improved interaction strength significantly to ∼57% and ∼67%, respectively (Fig. 5 e,f). Thus, the two mutations in the N-terminus of Pf-Ubc9 acted synergistically to facilitate interaction with Hs-SUMO1. More importantly, these mutants resembling Hs-Ubc9 showed an expected but mild reduction of ∼20-25% in their Pf-SUMO interaction strengths, comparing well with Pf-SUMO:Hs-Ubc9 interaction (Fig. 5 g,h).

### Pf-SUMO functionally engages the cellular human SUMOylation machinery

We asked if Pf-SUMO can utilize cellular SUMOylation machinery and modify host proteins inside cells. First, we checked the localization of Pf-SUMO by transfecting Venus-tagged Pf-SUMO into HeLa cells. Venus-tagged Hs-SUMO1 and vector alone served as the positive and the negative controls, Pf-SUMO and Hs-SUMO1 expression levels were comparable, and they presented an overlapping intracellular localization pattern with a significant signal inside the nucleus (Fig. 6 a). Second, to assess SUMOylation efficiencies, we enriched Venus-tagged SUMO from HEK293T cell lysates coexpressing Hs-Ubc9. We estimated Hs-SUMO1 modified proteins intensity (bands appearing above 50 kDa) to be 100% modification under these experimental conditions and found that Pf-SUMO-mediated modification was ∼30% (Fig. 6 b,c).

**Fig. 6:**
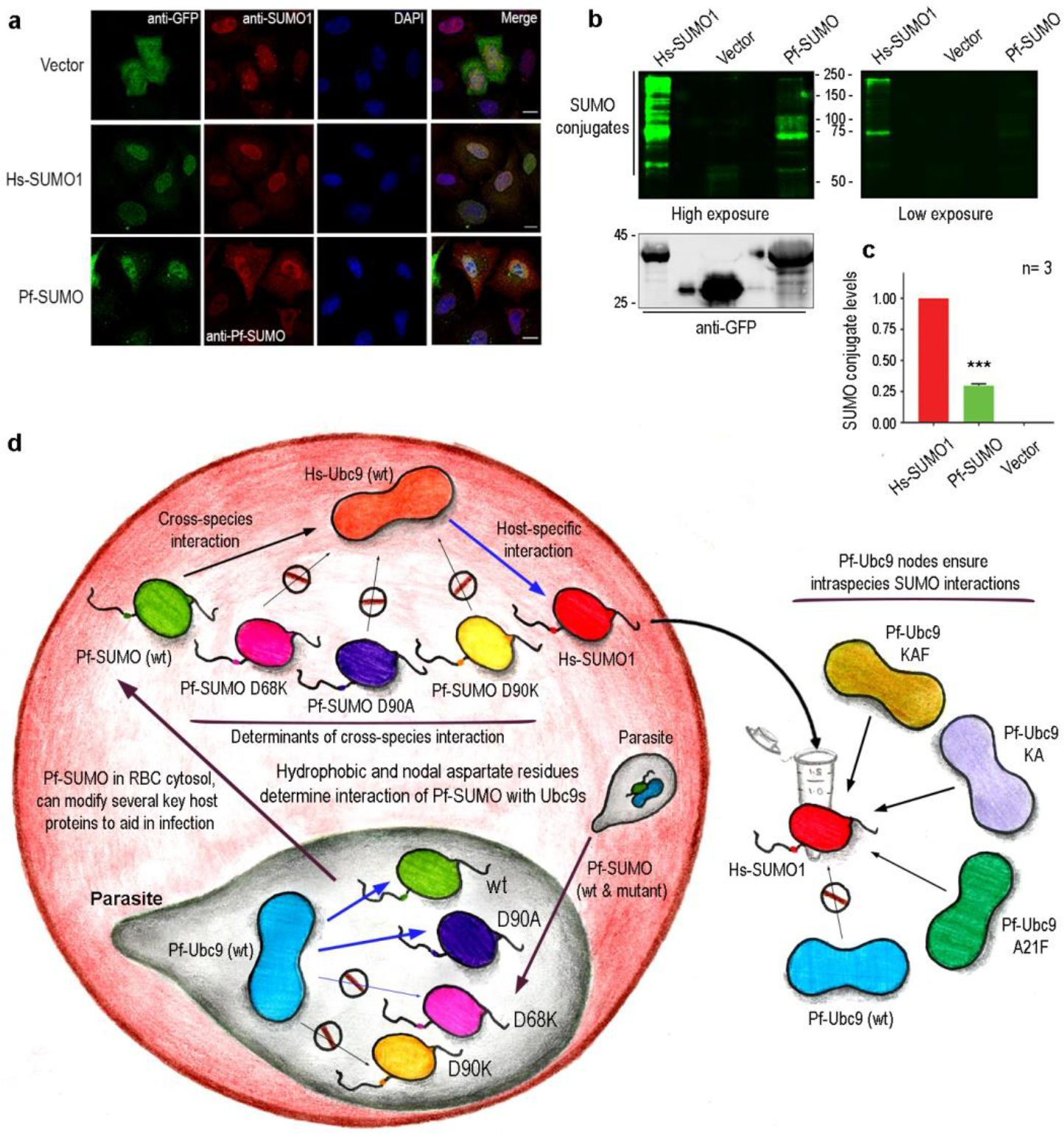
Pf-SUMO can utilize human SUMO machinery *in cellulo* to modify host proteins. **a)** Localization of Venus tagged Hs-SUMO1 and Pf-SUMO in Hela cells. Cells stained with anti-GFP antibodies (first vertical panels, green) and anti-Hs-SUMO1 or anti-Pf-SUMO specific antibodies (second vertical panels, red) under respective transfection conditions. Chromatin is visualized by DAPI staining (third vertical panels, blue). The scale bar represents 10µm in each case. **b)** GFP binding protein (GBP) mediated pulldown from HEK293T cell lysates expressing vector control, or Venus tagged Hs-SUMO1 or Pf-SUMO. The extent of SUMO conjugation in each case is assessed by western blotting with anti-GFP antibodies (left and right upper panels). Lower left panels indicate input expression levels of expressed proteins. **c)** The quantification of the relative *in cellulo* SUMO conjugation as seen in b. **d)** A working model for Pf-SUMO cross-reactivity with human Ubc9 enzyme. The model also elaborates the anticipated effects of nodal mutation in Pf-SUMO on its ability to interact with the human Ubc9 enzyme. It also suggests a possible role for Pf-SUMO in host RBC cytosol to modulate host responses by conjugating several host proteins. All experiments are performed at least three times independently. All statistical analysis was carried out using GraphPad Prism 8.4.3. Column analysis of data sets carried out by One-way ANOVA (nonparametric). Dunnett’s test was used for multiple comparisons. Family-wise significance and confidence level is p<0.001.

Next, we analyzed SUMO modified proteins enriched from Hs-SUMO or Pf-SUMO overexpressing HEK293T cells using mass spectroscopy. We identified several proteins pulled down from Hs-SUMO1 and Pf-SUMO transfected cells and assessed ∼60 proteins to be significant. We expected so and found that a significantly large number of these modified proteins were common between Hs-SUMO1 and Pf-SUMO. These SUMO-modified proteins are components of established cellular machinery and pathways like transcription regulation, stress response, protein chaperone, cytoskeletal components and regulators, and cellular signaling mediators(Supplementary Table 3). The MS analysis of Pf-SUMO modified host cell protein established the notion that Pf-SUMO can utilize host SUMOylation machinery, and the same can play an essential role in the sustenance of *Plasmodium* infection.

The model in Fig. 6 d summarizes and suggests that Pf-SUMO can interact with human SUMOylation machinery, and the C-terminal aspartates at 68^th^ and 90^th^ positions serve as critical nodes for discriminating Ubc9 enzymes leading to functional and opportunistic interaction. Moreover, Pf-Ubc9 has acquired critical changes in the N-terminal region allow only the proteins of the cognate SUMO pathway to interact. Thus, these residues serve as critical nodes in ensuring unidirectionality of cross-talk between Plasmodium and human SUMO pathway interactions. Selectively disrupting the interaction interface at the critical modes will offer an effective strategy against *Plasmodium* infection.

## Discussion

Covalent protein modification during PTMs is often governed by non-covalent (electrostatic, π−π hydrophobic) protein-protein interactions. SUMOylation is known to modulate protein-protein interactions, localization, and activity of the target proteins inside cells. Several pathogens target hosts SUMOylation pathway E1 and E2 enzymes to bolster infection and pathogenesis^22, 23, 31, 32^. Interestingly, the SUMOylation pathway specificity in many eukaryotes and *Plasmodium* lies at the E1-E2 enzyme interaction level. Sequential and simultaneous interactions exist among the E1, E2 enzymes, and the SUMO paralogs^13, 14, 16, 33–35^. Significant sequence similarities between *Plasmodium* and human SUMOs and Ubc9s, and a positive Pf-SUMO:Hs-Ubc9 interaction, undetectable Hs-SUMO1:Pf-Ubc9 interaction (Figure S1 and Figure 1) are important observations. Striking structural similarity between Pf-SUMO and Hs-SUMOs, and SUMO:Ubc9 interaction interface further supports the idea of molecular piracy by Plasmodium and argues favourably for a physiologically relevant presence of Pf-SUMO’s in the RBC cytosol^9^.

Charge-based non-covalent interactions between SUMO C-terminus and Ubc9 N-terminus are critical for SUMO modification^18, 19, 33, 34^. Studies highlight the importance of negatively charged residues in SUMO/Smt3 C-terminus^30^. Mutation of D68 residue in Pf-SUMO to alanine or lysine abolished interaction with Pf-Ubc9 and Hs-Ubc9 and a cascading effect in its ability to participate in SUMOylation (Fig. 4). We propose that the D68 of Pf-SUMO is equivalent to E67 in Hs-SUMO1. E67 is an integral part of the E67 interacting loop (EIL), exhibits precise non-covalent interactions^36, 37^. Further, D68 of Pf-SUMO can play defining role in RBC cytosol to help in malaria pathogenesis. The D68 residue perhaps can be part of a canonical <u>p</u>rotein <u>ex</u>port <u>el</u>ement (PEXEL) motif^38–42^, aiding in the possible translocation of Pf-SUMO into the RBC cytosol to exert effects on cross-species interaction with Hs-Ubc9.

Contrasting observations made with the Pf-SUMO 90^th^ position aspartate mutants, where Pf-SUMO D90A: Hs-Ubc9 interaction decreases, with negligible impact on Pf-SUMO D90A:Pf-Ubc9 interaction (Fig. 4) suggested a discriminating role for the D90 residue in Ubc9 specificity selection. Careful analysis of the Pf-SUMO D90A:Pf-Ubc9 interaction indicated that the charge neutralization should induce loss of a salt bridge formation, but phenylalanine (F23) of Ubc9 and alanine (A90) of Pf-SUMO D90A mutant helps in restoring interaction through a new compensatory hydrophobic interaction (Fig. 5). These results also help explain the reason for the lack of cross-species interaction between Hs-SUMO1:PfUbc9 (Fig. 1). Further studies analyzing the anchor points will help assert a regulation on cross-species interaction between Pf-SUMO and Hs-Ubc9.

The N-terminal region of Ubc9 determines the species-specificity by engaging in specific interactions individually with E1 and SUMO. Accordingly, complete substitutions of Pf-Ubc9 N-terminus (aa 1-81) with residues of Hs-Ubc9 allowed functional interaction with the human-E1 enzyme^10, 11^. Similarly, we report Alanine 13, Glutamate 14, and Alanine 21 in the N-terminus of Pf-Ubc9 as specific nodes determining cross-species interaction with Hs-SUMO1 (Fig. 5). Thus, these alterations in the N-terminus of Pf-Ubc9 appear an exciting strategy in engaging species-specific and cross-species interactions.

Our analysis centered around D68 and D90 of Pf-SUMO align well with synthetic lethality and growth defect phenotypes of equivalent charge-reversal mutations in the yeast Smt3^30^. Thus, we propose that D68 and D90 of Pf-SUMO as critical nodes in establishing the cross-species and species-specific interaction with Ubc9s. Further, it can be extrapolated that D90 residue is more critical for engaging in species-specific Ubc9 recognition, differentiating Pf-Ubc9 and Hs-Ubc9 enzymes. Likewise, the N-terminal region of Pf-Ubc9 is a critical determinant of species-specific interaction with Hs-SUMO and may provide a handle to the parasite to exploit the host machinery (Fig. 6 d).

In a host-pathogen interaction scenario, exploiting the host’s pathways with the most negligible impact on its pathways represents an expedient strategy of a successful pathogen. Pf-SUMO’s presence in Maurer’s Cleft may be a similar strategy, suggesting Pf-SUMO can utilize the host SUMOylation machinery and modify essential host proteins. We propose that *Plasmodium*, through Pf-SUMO, can modulate the host processes and can exploit the human SUMOylation pathway during intraerythrocytic development stages. Besides, Hs-SUMO1 and Pf-Ubc9 interactions do not make sense in evolutionary terms. Several pathogens target the Ubc9 enzyme of the host and affect the SUMOylation process. We demonstrate that Pf-SUMO achieves the same while staging molecular piracy of the human SUMOylation pathway for successful infection. Importantly, targeting of SUMOylation pathway to regulate *Plasmodium* infection^43^ can be an efficient strategy. Interestingly, small molecule inhibitors generated against *Plasmodium* SENP1 hold strong potential for malaria therapy. In the future, the SUMOylation pathway and the differentiating interface between SUMO and Ubc9 in host and pathogen can be exploited to target selective interface inhibitors as a possible remedy against malaria pathogenesis.

## Methods

### cDNA Cloning and Plasmid Construction

The mature form of *Pf*-SUMO (*Pf*-SUMO-GG) and *Pf*-Ubc9 enzyme coding regions were PCR-amplified from λZAP Plasmodium genomic DNA library provided by Prof. Shobhona Sharma (TIFR Mumbai, India) and subsequently cloned into pGEX-6P-1. The pGEX-6P-1 constructs thus generated have a GST-tag at the N-terminal end followed by the PreScission protease recognition site. Further, the pGEX-6P-1 clones of Pf-SUMO and Pf-Ubc9 were received from Prof. Michael Matunis (Johns Hopkins Univ. Baltimore, USA). *Hs*-SUMO1 and *Hs*-E2 were also sub-cloned into a pGEX-6P-1 vector. The wildtype and *Pf*-SUMO mutants described earlier were PCR amplified and subcloned into mammalian expression vector pVenus-C1. Human and Plasmodium Ubc9 were subcloned into the pET-28a (+) vector for expression and purification of (His)_6_-tagged recombinant proteins. The *Pf*-SUMO wild type clone (pSUMO1-1S) for *in bacto* SUMOylation was derivatized from pSUMO1, was a gift from Primo Schaer (Department of Biomedicine, University of Basel; Addgene plasmid #52258). All *Pf*-SUMO and *Pf*-Ubc9 mutants were generated using Q5 Site-Directed Mutagenesis kit (NEB #E0554S) in *Pf*-SUMO wild type backbone. The chimera of *Pf*-SUMO and human SUMO-1 were generated by utilizing the internal EcoRI site in the cDNA of *Hs*-SUMO1. For PfN chimera, the N-terminal coding region of *Pf*-SUMO (aa 1-60) was PCR amplified with BamHI and EcoRI ends and ligated pGEX-6P-1-*Hs*-SUMO1 clone digested to generate the compatible ends. To generate PfC chimera, the C-terminal coding region of *Pf*-SUMO (aa 61-98) was PCR amplified with EcoRI and XhoI ends and was ligated into pGEX-6P-1-*Hs*-SUMO1 clone digested to generate the compatible ends. The veracity of all the constructs used in the study was established by DNA sequencing.

### Recombinant Protein Expression and Purification

Unlabeled and isotope-labeled proteins were all expressed in *E.coli* BL21 (λDE3) cells. All proteins such as Hs-SUMO1, Hs-E2, Pf-E2, and Pf-SUMO (WT and its mutants) were expressed and purified as previously described^44^. Briefly, cell pellets were resuspended in lysis buffer (20 mM Tris-HCl pH 8.0, 1 mM EDTA, 100 mM NaCl) containing 0.01% Triton X-100, 1 mM PMSF (phenyl methane sulfonyl fluoride) protease inhibitor and 1 mg/mL of lysozyme. Resuspended cell pellets were sonicated (30% amp, pulse on 3 sec, pulse off 5 sec) for 20 min in ice and centrifuged at 17,000 rpm for 45 min at 4 °C to pellet down the cell debris. The supernatants were incubated for 2 h at 4 °C with Glutathione-Agarose beads to bind GST-tag proteins to the beads. Beads were washed with lysis buffer containing increasing NaCl concentration (200, 400, and 600 mM) to remove non-specifically binding proteins. *Pf*-SUMO (WT) and its mutants and Pf-E2 proteins were digested with PreScission protease on-column cleavage at 4 °C. However, Hs-SUMO1 and Hs-E2 proteins were digested with thrombin (3-5 units/mg of proteins) at 22 °C to remove the GST tag from the GST-tag proteins. All eluted proteins were further purified on a Superdex^TM^ 75 10/300 GL column (GE Healthcare). The purified proteins were analyzed by SDS-PAGE and MALDI-TOF and concentrated using ultra-filtration through a 3 kDa cut-off Amicon (Millipore) membrane. The concentration of proteins was estimated using UV-absorption at 280 nm. For NMR experiments, singly labeled (^15^N) and doubly labeled (^13^C and ^15^N) protein samples were prepared using ^15^NH_4_Cl and uniformly ^13^C labeled glucose as the sole sources of nitrogen and carbon. For isothermal titration calorimetry (ITC) and SPR analysis, eluted proteins were dialyzed into a buffer containing 20 mM Tris-HCl, pH 7.8, 50 mM NaCl.

For *in vitro* pulldown assays and SUMOylation experiments, proteins were expressed in *E.coli* BL21 (λDE3) cells and induced with 200 µM isopropyl β-D-1-thiogalactopyranoside (IPTG) for 4 h at 30 °C. For GST tagged protein purification, cell pellets were resuspended with lysis buffer (20 mM Tris-HCl pH 7.5, 1 mM EDTA, 150 mM NaCl) containing 0.1% Triton X-100, 5 mM β mercaptoethanol (β-ME), 1 mM PMSF (phenyl methane sulfonyl fluoride, MP Biomed #195381) protease inhibitor and 1 mg/mL of lysozyme. Resuspended cell pellets were sonicated (45% amp, pulse on 10 sec, pulse off 20 sec) for 3 min on ice and centrifuged at 20,000 rpm for 30 min at 4 °C to pellet down the cell debris. The supernatants were incubated for 1 hr at 4 °C with Glutathione-Agarose beads to bind GST-tag proteins to the beads. (Merck #70541). Beads were washed with lysis buffer containing 400 mM of NaCl to remove the non-specific binding of other proteins. Proteins were eluted using 20 mM glutathione-containing lysis buffer (MP Biomed #101814). All eluted proteins were further dialyzed using a 12-14 kDa dialysis membrane (Spectra/Por #132706) to remove excess glutathione. For (His)_6_-tagged protein purification, cell pellets were resuspended with lysis buffer (50 mM NaH_2_PO_4_ pH 8.0, 300 mM NaCl, 10 mM imidazole [MP Biomed #102033]) containing 0.1% Triton X-100, 5 mM β mercaptoethanol (β-ME), 1 mM PMSF protease inhibitor and 1 mg/mL of lysozyme. Resuspended cell pellets were sonicated (45% amp, pulse on 10 sec, pulse off 20 sec) for 3 min in ice and centrifuged at 20,000 rpm for 30 min at 4 °C to pellet down the cell debris. The supernatants were incubated for 1 h at 4 °C with Ni-NTA agarose beads to bind (His)_6_-tag proteins to the beads. (Qiagen #1018244). Beads were washed with lysis buffer containing 25 mM of imidazole to remove the non-specific binding of other proteins. Proteins were eluted using the lysis buffer containing 250 mM imidazole (MP Biomed #101814). All eluted proteins were further dialyzed using 12-14 kDa dialysis membrane to remove the imidazole. Further, proteins were quantitated and stored at -80 °C as aliquots.

### Antibody generation

Antibodies against Pf-SUMO were custom generated at Imgenex, India. Bacterially purified GST-Pf-SUMO were injected into rabbits, and polyclonal antibodies were affinity purified over GST-Pf-SUMO cross-linked beads. Purified antibodies were characterized for their efficacies in western blotting and immunofluorescence experiments. In addition, antibodies for Human-SUMO1 were generated as described elsewhere^45^.

### Structure calculations using NMR spectroscopy

Isotopically labeled proteins for NMR experiments were prepared as published in protocol elsewhere^44^. For structure calculation, samples of ^15^N and ^13^C/^15^N-labelled *Pf*-SUMO were prepared at a concentration of 1.0 mM in 20 mM Tris-HCl pH 7.0, 50 mM NaCl, containing 90% H_2_O, 10% ^2^H_2_O. All the NMR experiments were performed on Bruker (AVANCEIII HD) 750 MHz spectrometers equipped with a room temperature triple resonance probe equipped with Z-gradient. A series of two- and three-dimensional experiments like ^15^N-edited (τ_mix_-150 ms) and ^13^C-edited NOESY-HSQC (τ_mix_-150 ms) spectra were recorded to generate the structural restraints for *Pf*-SUMO protein at 298 K. All spectra were processed using Bruker TOPSPIN 3.2, and NOE cross-peaks were assigned manually using CARA 1.8.4.2 software^46^. Intensity obtained from cross-peaks was used to generate distance restraints using the CYANA-3.0^47^ program. Out of 1415 distance restraints, the distribution of various NOE restraints was as follows: 245- sequential, 367- intra-residual, 342,- medium-range and 461- long-range. In addition, 40 hydrogen bonds obtained for H-D exchange ^15^N-HSQC experiments were also used as restraints. CYANA 3.0 software was used to generate 200 randomized conformers, out of which 10 conformers with the lowest target function, having no distance and angle violation, were selected. These 10 conformers were further refined using the CNS 1.21 software based on molecular dynamics simulation and the standard water shell refinement protocol^48, 49^. The distances between the atoms are relaxed during this stage, improving Ramachandran’s plot statistics and Z-score for the (phi, psi) residues in the ordered region. The PSVSv1.4 software (http://www.psvs-1_4.nesg.org) was used to analyze the quality of the structure. PYMOL software (http://pymol.sourceforge.net/) was used for generating figures for structures.

### Isothermal Titration Calorimetry

ITC experiments were performed for Pf-SUMO, D90A-Pf-SUMO, and Hs-SUMO1 with Pf-Ubc9 and Hs-Ubc9 using MicroCal iTC200 (GE Healthcare) in their respective buffer. Pf-SUMO and its mutant and Hs-SUMO1 (2 µL, 450-600 μM) were added with the syringe to the sample cell containing (30-50 μM) Pf-Ubc9 and Hs-Ubc9 at a constant stirring rate of 1000 rpm. A total of 19 injections was performed for each experiment with an interval of 120 sec. In each injection, the mixing duration between cell and syringe sample was 5.0 sec. To nullify the heat of dilution, Pf-SUMO and Hs-SUMO1 were titrated against a buffer and subtracted from the raw data prior to model fitting. The temperature was maintained at 298K during the experiments. Titrations were performed in duplicate using the same set of stock solutions. The ITC data were analyzed using the ORIGIN version of the software provided by MicroCal iTC200.

### Surface Plasmon Resonance

The binding kinetics of Pf-SUMO, D90A-Pf-SUMO, and Hs-SUMO1 with Pf-Ubc9 and Hs-Ubc9 were determined using surface plasmon resonance (BIAcore T200 GE Healthcare) at 298K. The Pf-SUMO, D90A-Pf-SUMO, and Hs-SUMO1 proteins were immobilized in 10 mM sodium acetate buffer (pH 4.5) on the CM5 sensor chip. The various concentrations (0.78-100 µM) of Pf-Ubc9 and Hs-Ubc9 were passed over the immobilized proteins at a flow rate of 30 µL/min. The 20 mM Tris-HCl, 50 mM NaCl (pH 7.8) was used as a running buffer. The contact time and dissociation time were 120 and 300 sec, respectively. For regeneration, 10 mM Glycine (pH 2.5) was used. The obtained sensorgram is fitted into a steady-state affinity equation using Biacore T200 evaluation software.

### NMR titrations of Pf-SUMO and its mutant against Pf-Ubc9 and Hs-Ubc9

To check the interactions of Pf-SUMO with Pf-Ubc9 and Hs-Ubc9, we recorded a series of HSQC spectra of the ^15^N labeled Pf-SUMO and its mutant in its free form and then titrated with a different equivalent of unlabelled Pf-Ubc9 and Hs-Ubc9 enzymes (0.05, 0.1, 0.2, 0.4, 0.6 equivalents). Similar experiments were performed with ^15^N labeled Hs-SUMO-1 and different equivalents of unlabelled Pf-Ubc9 and Hs-Ubc9 enzymes. Perturbation in amide cross peak either due to decreased intensity or chemical shift perturbation (CSP) was monitored. CSP was calculated by using the formula ΔΔδ = [(5ΔδH^N)^)^2^ + (Δδ^15^N)^2^]^1/2^ where δH^N^ and δ^15^N represent the difference in proton and nitrogen chemical shifts respectively. Similarly, intensity change was quantified as the amide cross peaks intensities (I) with respect to the same cross-peaks intensities (I_0_) in the absence of E2 proteins.

### Docking study of Pf-SUMO and Hs-SUMO1 against Pf-Ubc9 and Hs-Ubc9 enzymes

The crystal structure of Hs-SUMO1 with Hs-Ubc9 (PDB code: 2uyz) was taken as a template to build the docked model for Pf SUMO with Pf-Ubc9 and Hs-Ubc9 enzymes. We used the HADDOCK (High Ambiguity Driven, protein-protein Docking) server 2.2 (http://haddock.science.uu.nl/services/HADDOCK2.2) for protein-protein docking^50^. HADDOCK uses NMR restraints as an input parameter to perform a guided docking. Here Pf-Ubc9 and Hs-Ubc9 were taken as a ligand, and the NMR structure of Pf-SUMO ((PDB code: 5gjl) was taken as a receptor. We provided the active and passive residues originated from the NMR titration study for docking. The detailed information about the interface area of docked models was analyzed by using the web-based server PDBePISA (http://www.ebi.ac.uk/msd-srv/prot_int/pistart.html).

### *In vitro* SUMOylation reaction

*In vitro* SUMOylation reactions contained E1 enzyme (0.25 µg GST-SAE2/SAE1), E2 enzyme (1.0 μg (His)_6_-Ubc9), GST-tagged SUMO protein (4 μg of wildtype or mutant SUMO), and GST-tagged substrate peptide (4 μg) in the reaction buffer (50 mM Tris, pH 7.5, 5 mM MgCl_2_, 5 mM ATP, 5 mM DTT). SUMOylation reactions were incubated at 37 °C for indicated time points and terminated with 6X Laemmlli buffer and boiled for 10 min. SUMOylation reaction was analyzed by resolving reaction products on SDS-PAGE followed by Coomassie Brilliant Blue (R250) staining. Quantification of *in vitro* SUMOylation was carried out at an interval of 3 h.

### *In bacto* SUMOylation reaction

A standard substrate peptide constructs in pGEX-6P-1 vector co-transformed with pSUMO1-1S (Pf-SUMO wildtype or its mutants) to *E.coli* BL21 (λDE3) cells. Double transformants were selected on Luria broth (LB) agar plates having 50 mg/L of ampicillin and 25 mg/L of streptomycin. For *in bacto* SUMOylation, mid-log phase cultures were induced with 200 µM isopropyl β-D-1-thiogalactopyranoside (IPTG) at 25 °C for 18 h. Cells were harvested by centrifugation at 10,000xg, lysed in lysis buffer (20 mM Tris, pH 7.5, 150 mM NaCl, 1 mM EDTA, 0.1% Triton X-100, 5 mM β-ME, 1 mM PMSF) and soluble protein fractions were extracted by sonication. Crude lysates were then cleared by centrifugation at 20,000 rpm at 4 °C for 30 min. Recombinant proteins were affinity purified on Glutathione-Agarose beads followed by washes with wash buffer containing 400 mM NaCl. For visualization of bead-bound purified proteins, beads were boiled in 1X Laemmli buffer, and the resultant sample was subsequently analyzed on SDS–PAGE followed by Coomassie Brilliant Blue (R250) staining.

### *In vitro* pulldown reaction

For *in vitro* pulldown experiments, 13 μM (His)_6_-tagged Ubc9 were immobilized on the Ni-NTA beads, and 20 μM of purified wildtype or mutant SUMOs were mixed in binding buffer (20 mM Tris HCl pH 7.5, 150 mM NaCl). Tubes were incubated under constant rotation (300 rpm) at room temperature for 2 h. Subsequently, beads were washed three times carefully with wash buffer (20 mM Tris HCl pH 7.5, 150 mM NaCl) for 5 min each. Finally, bead-bound proteins were extracted by boiling for 10 min in 1X Laemmli buffer. Samples thus obtained were further analyzed on SDS-PAGE and western blotting.

### Western Blotting

All samples for western blotting were resolved to the desired extent on SDS-PAGE. Following wet transfer protocols, proteins were transferred onto the methanol-activated µm PVDF membrane (Merck #ISEQ85R) using 1X transfer buffer (2.5 mM Tris-HCl pH 7.5, 19.2 mM Glycine). The membrane was blocked for 1 h in 5% BSA. Further, the membrane was incubated overnight with primary antibodies (Rabbit-anti-Hs-SUMO1 [1:5000], or Rabbit-anti-Pf-SUMO [1:5000]). The membrane was later washed three times for 10 min each with TBS-T buffer (20 mM Tris HCl pH 7.5, 150 mM NaCl, 0.1% Tween-20). The membrane was incubated with Alexa fluor Plus 680 secondary antibodies (Invitrogen #A32734) for 1 h. Later it was washed with TBS-T buffer (three times for 10 min each). Images were taken using LI-COR (Model: 9120) IR system.

### Cell culture and transfections

HEK293T and HeLa cell lines were cultured in Dulbecco’s Modified Eagle’s Medium (DMEM) (Gibco #11995-065) supplemented with 10% (v/v) Fetal Bovine Serum (FBS) (Gibco #10270-106) and 1% antibiotics (Gibco #15240-062) in a humidified incubator at 37°C under 5% CO_2_ conditions. 1 x 10^7^ cells were grown in 100-mm cell culture plates for GFP-trap pull-downs or on coverslips in a six-well plate format (3 x 10^5^ cells per well) for immunofluorescence. 12 hrs post-seeding, cells were transfected using polyethylenimine (PEI) 25-kDa linear polymer (Polysciences Corporation Ltd. #23966) or Effectene Transfection reagent (Qiagen #301425) following the manufacturer’s instructions.

### Immunofluorescence

HeLa cells were transfected with desired mammalian SUMO expression clones using Effectene Transfection reagent and allowed to grow for 12 hrs. Later, cells were again washed twice with PBS and fixed using 4% formaldehyde for 15 min at 4 °C. The cells were then rehydrated and permeabilized with rehydration buffer (10 mM Tris, 150 mM NaCl, 0.1% TritonX-100) for 10 min. Cells were blocked with 5% Normal Goat Serum (NGS) for 1 h at 4 °C after rehydration. The cells were stained overnight at 4 °C with rabbit-anti-Hs-SUMO1 (1:600), rabbit-anti-Pf-SUMO (1:600), and the anti-GFP antibody (1:800, sc-9996, Santa Cruz Biotechnology). After primary antibody incubation, cells were washed three times with PBS-T (5 min each) and incubated with 1:1000 dilutions of anti-mouse Alexa Flour 488 (Thermo #A11029) and anti-rabbit Alexa Fluor 568 (Thermo #A11036) for 1 h. Cells were washed thrice with PBS-T and mounted on slides using DAPI containing mounting medium (Sigma #F6057). Fluorescence signals were captured on Zeiss LSM 780 confocal microscope, and images were analyzed using Image J software.

### GFP-trap pulldown

GFP binding protein (GBP) clone was a kind gift from Heinrich Leonhardt (Ludwig-Maximilians-University of Munich). GBP cDNA was subcloned into the pGEX-6P-1 vector to express GST-tagged GBP. Venus tagged Hs-SUMO1 and Venus-tagged Pf-SUMO were coexpressed with 3X-FLAG-Ubc9 (100-mm dish format) in HEK293T cells. Cleared lysates prepared from cells expressing GFP-SUMO variants and were mixed with GST-GBP, and the complex formed was pulled down on glutathione-agarose beads. Bound material was eluted and processed in 1X Laemmli buffer for Western blotting and detected with anti-GFP antibody (1:6000, sc-9996, Santa Cruz Biotechnology).

### Mass spectrometry analysis of GFP-trap pulldown proteins

Mammalian proteins enriched by GFP-trap pulldown were subjected to mass spectrometry analysis. SCIEX X500B qToF platform paired with ExionLC AD UHPLC and XB-C18 column was used for the LC-ESI-MS data of trypsin digested proteins. The mass spectrometry data obtained were searched against the *Homo sapiens* database consisting of 20,395 proteins acquired from Uniprot using Proteome Discoverer 2.2 software. The processing workflow comprised of spectrum selector and SEQUEST incorporated as search engines. The search was carried out with trypsin, and double missed cleavage was allowed with a minimum peptide length of 6 amino acids. For MS1 peaks, the mass deviation was considered as 10 ppm and 0.05 Da for peptide tolerance. Carbamidomethylation of cysteine was included as static, and oxidation of methionine and acetylation were as dynamic modification. A false discovery rate (FDR) of 1% was considered for the result.

### Statistical analysis

Image analysis, processing, and band quantitation were done using ImageJ software or GelQuant.NET. The experiments were independently repeated at least three times, and the values are expressed as mean ± SD. Column analysis of data sets was carried out by One-way ANOVA (nonparametric) and Dunnett’s test for multiple comparisons. Family-wise significance and confidence level is p<0.001. Graphs were plotted using GraphPad Prism 8.4.3.

## Supplementary Figures and tables

**Supplementary Fig. 1:**
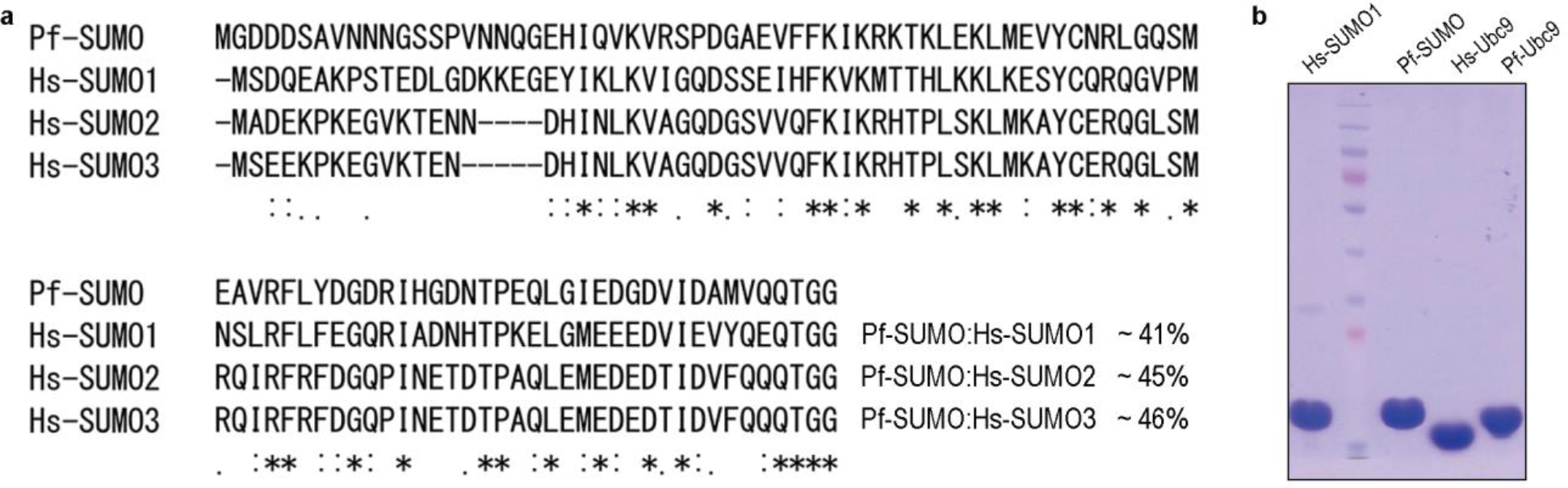
Protein purification and sequence alignment. **a)** Primary sequence alignment of Plasmodium SUMO against human SUMO1, SUMO2, and SUMO3. Percentage sequence identity of Plasmodium SUMO with each human SUMO paralog is mentioned. **b)** SDS-PAGE image showing untagged and purified Hs-SUMO1, Pf-SUMO, Hs-Ubc9, and Pf-Ubc9 proteins.

**Supplementary Fig. 2:**
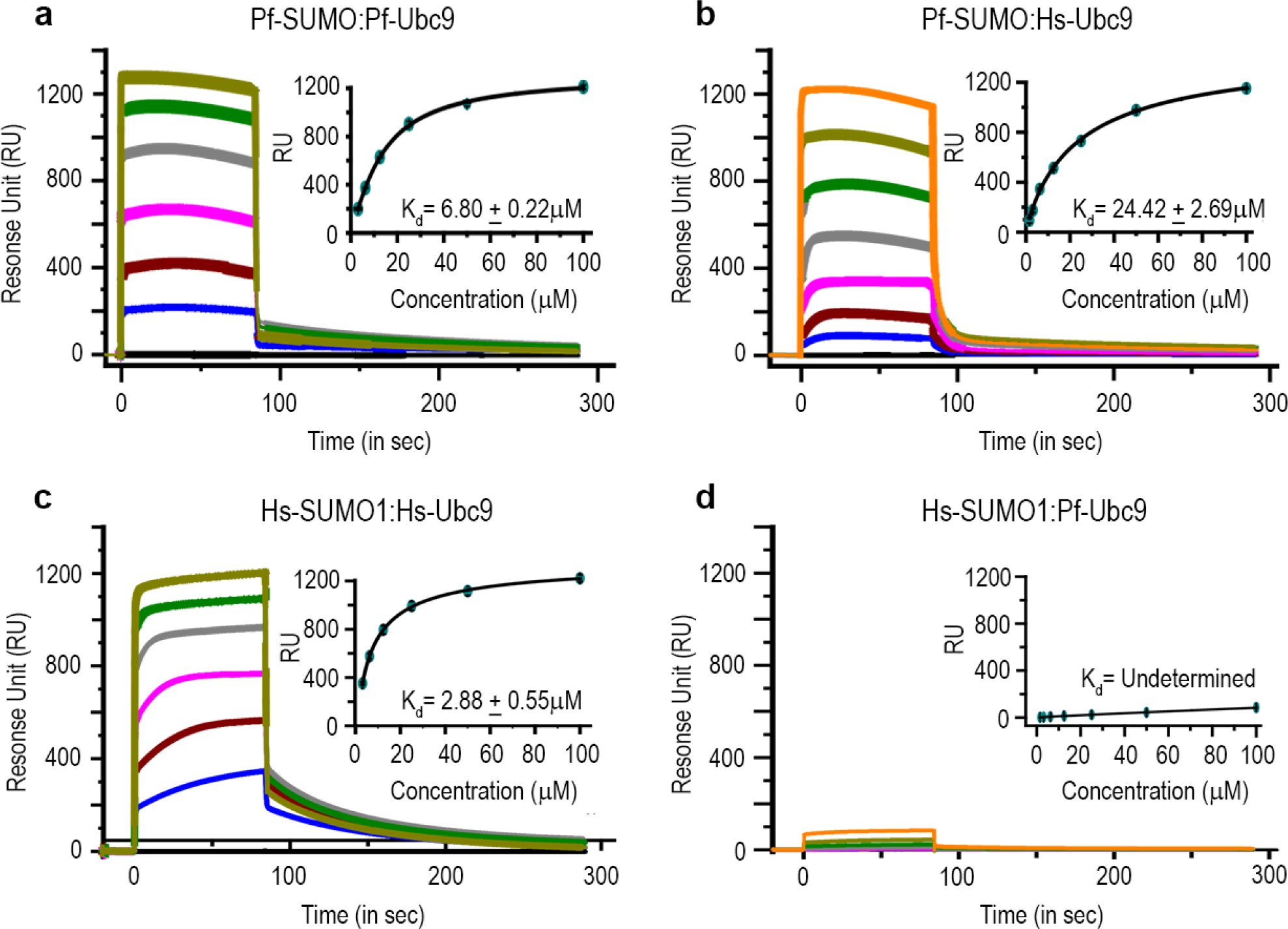
*Plasmodium* SUMO exhibits strong cross-reactivity with the human Ubc9 enzyme. Surface plasmon resonance-based binding studies for Pf-SUMO and Hs-SUMO1 interactions with Ubc9 enzymes. **(a and b)** The response unit of Pf-SUMO at different concentrations of Ubc9 enzyme and curve-fitting by 1:1 binding. **(c and d**) Same as in (a) and (b), but the Hs-SUMO1 was used instead of Pf-SUMO. The calculated binding affinities (K_d_) are indicated for each case. The interactions presented in (a) and (c) serve as a positive control in these SPR analyses.

**Supplementary Fig. 3:**
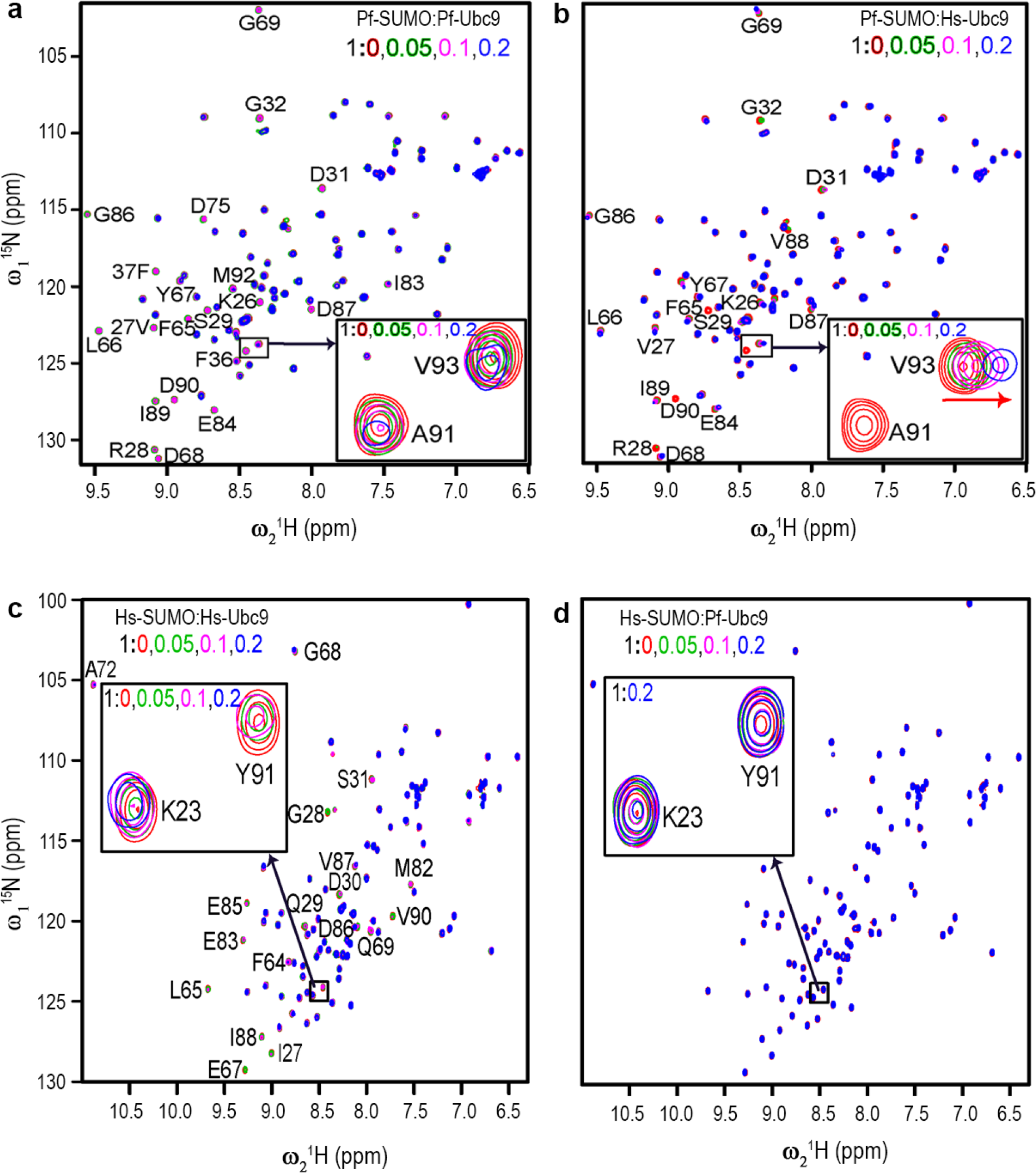
*Plasmodium* SUMO exhibits strong cross-reactivity with human Ubc9 enzyme. NMR-based titration experiments for Pf-SUMO and Hs-SUMO1 with Ubc9 enzymes. **(a and b)** Overlay ^15^N-^1^H heteronuclear single-quantum coherence (HSQC) spectra of Pf-SUMO with Pf-Ubc9 and Hs-Ubc9 enzymes respectively at a ratio of 1: 0.0, 0.05, 0.1, 0.2 i.e. free state (red colour) to the bound state (0.05, green; 0.1, magenta; 0.2, blue). The marked amino acid in HSQC spectra indicates the disappeared or shifted residues during the course of titration. The insets are excerpts of selected peaks from Pf-SUMO interactions as observed. The arrow in the inset indicates the direction of chemical shift perturbations. **(c and d**) Same as in **(a** and **b)**, but the Hs-SUMO1 was used instead of Pf-SUMO.

**Supplementary Fig. 4:**
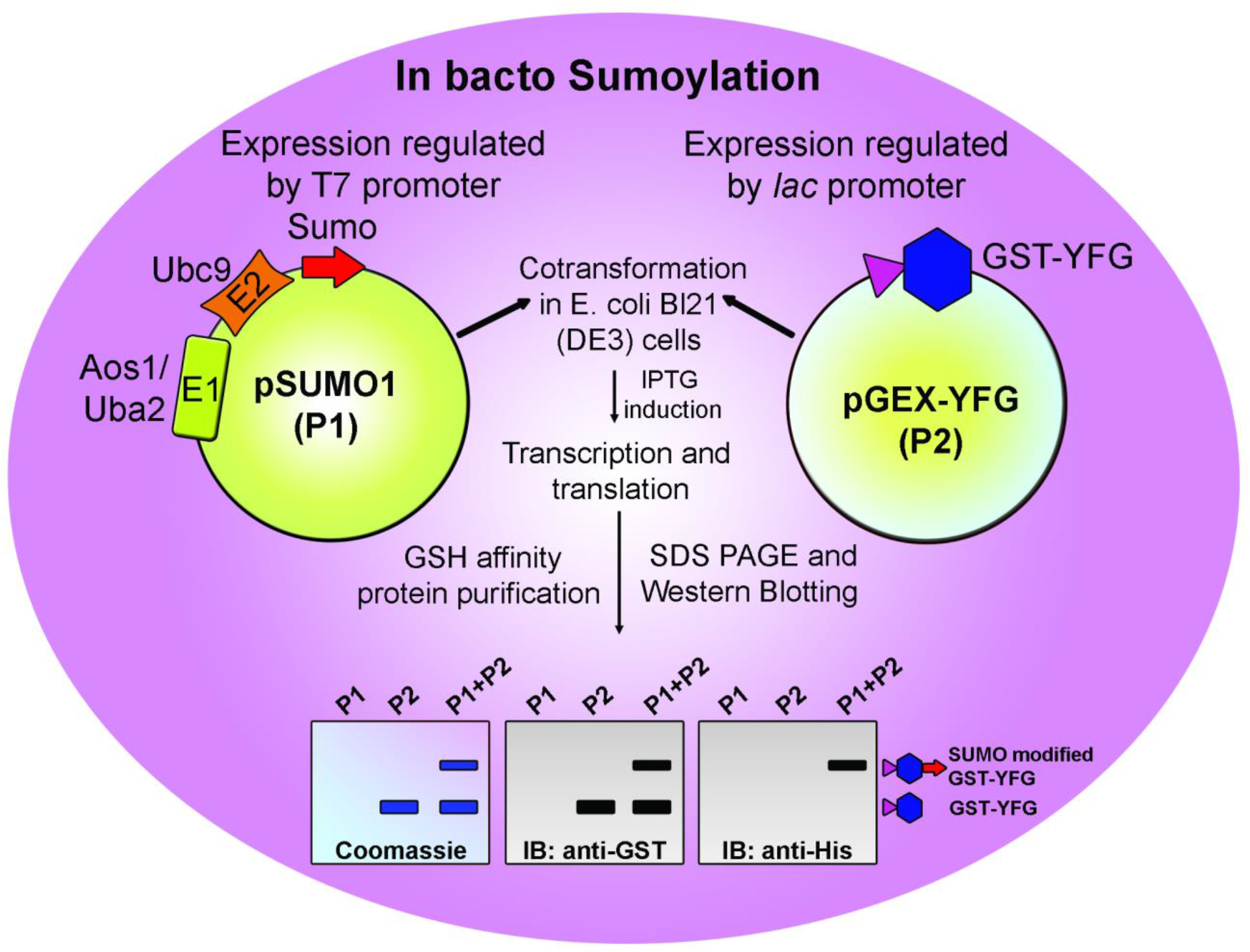
Illustration of *in bacto* SUMOylation. Bacterial expression plasmid vectors, P1 for expressing SUMO1 and E1 and E2 enzymes of the human SUMOylation pathway, and P2 for expressing GST tagged target protein (YFG) into *E. coli* Bl21 (DE3) cells. IPTG-based induction allows expression of SUMOylation pathway and target proteins. Glutathione (GSH) affinity purification enriches unmodified and SUMO-modified target protein from the total bacterial lysate. The possible SUMO modification of target protein can be assessed by slower migrating bands on coomassie stained SDS-PAGE gel. Further, the modified target protein can be detected by anti-GST (recognizing target protein forms) and anti-His (recognizing unconjugated SUMO1 and modified target proteins) antibody-mediated western blotting methods.

**Supplementary Fig. 5:**
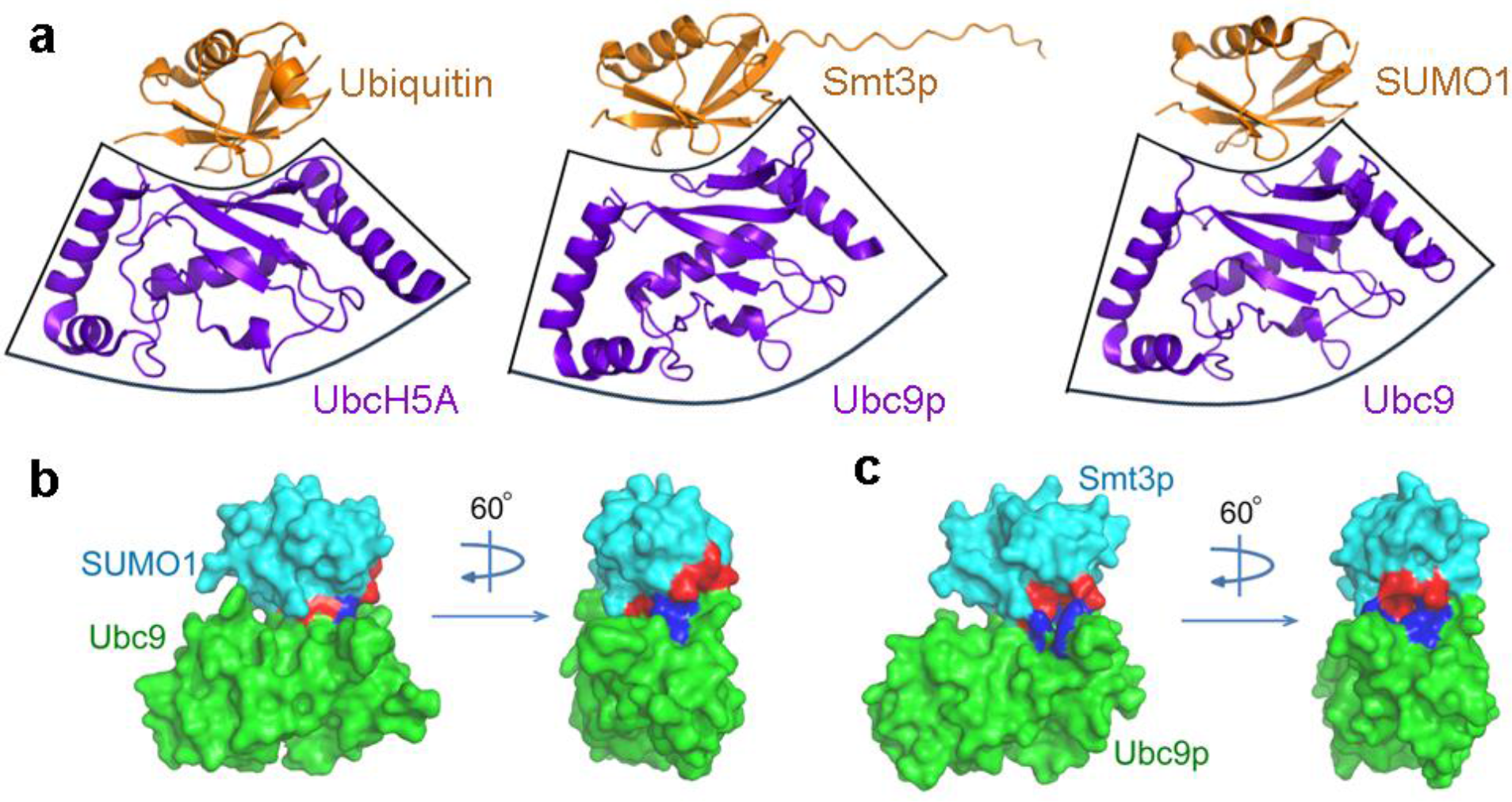
Shape and charged-based interaction between SUMO/Ubiquitin and their E2 counterpart. **a)** Structural comparison of the interaction between Ubiquitin and UbcH5A (PDB ID: 3PTF), human SUMO1 and human Ubc9 (PDB ID: 2PE6), and *Saccharomyces cerevisiae* ubiquitin-like protein Smt3p and Ubc9p (PDB ID: 2EKE) highlights the conserved concave-shaped interface between the ubiquitin-like proteins and Ubc9 across the species. Surface-filled structure of human SUMO1-Ubc9 complex in **(b),** and *S. cerevisiae* Smt3p-Ubc9p complex in (**c).** The structure has been rotated by 60**^◦^** to highlight the charged residues present at the interface. Interaction in both the complex is majorly driven by the electrostatic interaction between the positively charged residues, lysine in the Ubc, and negatively charged residues in the SUMO1 and Smt3p protein. Only the globular domain of Smt3p is represented in the surface-filled model (c) for a better representation of the charge-based interaction interface.

**Supplementary Fig. 6:**
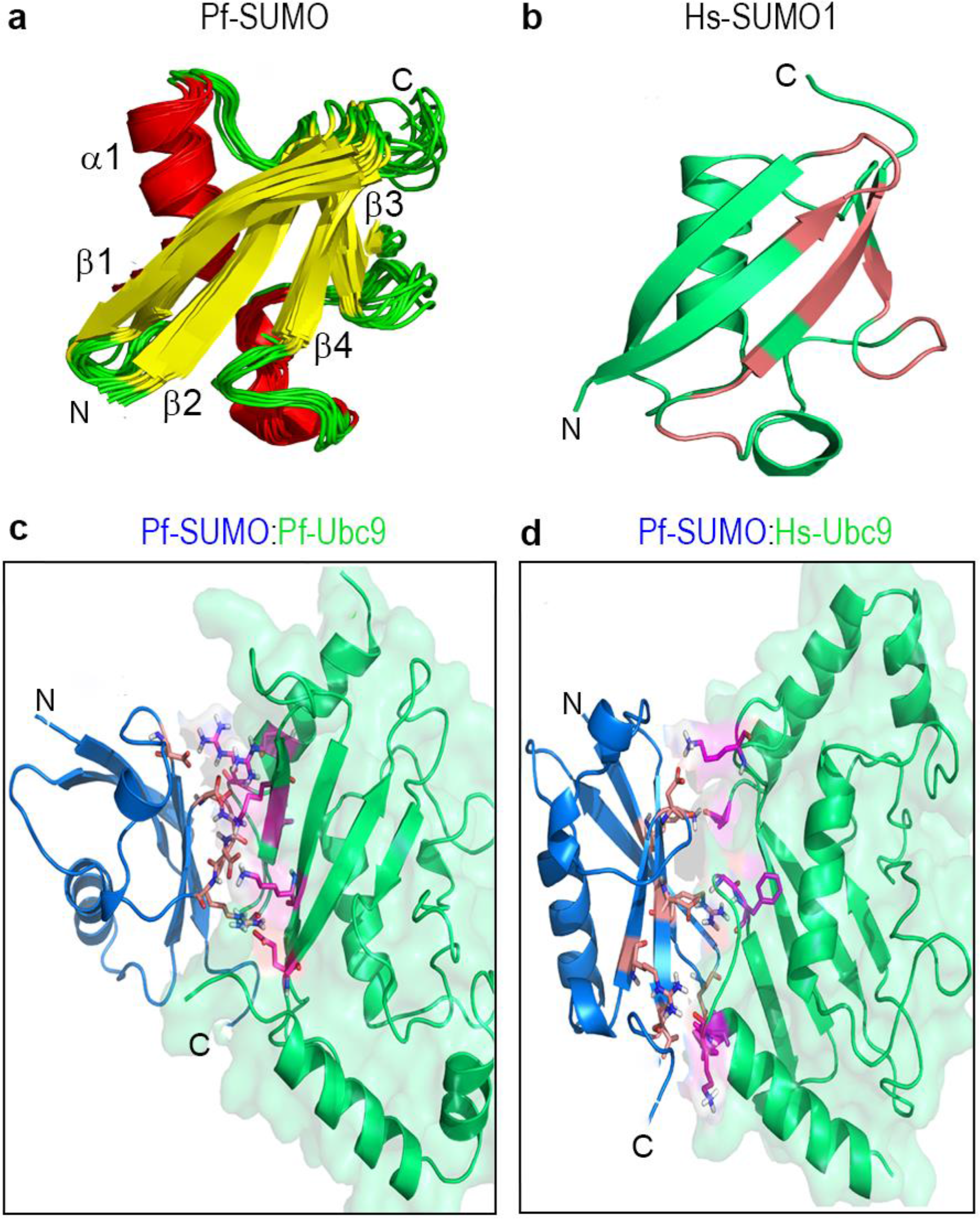
NMR-derived solution structure of Pf-SUMO and molecular docking of Pf-SUMO protein with E2 enzymes. **a)** Cartoon representation of the ten lowest energy structures of Pf-SUMO. The individual β-strands and α-helices are labeled. The β-strands are β1 (I23-V27), β2 (V35- I39), β3 (V63-L66), and β4 (D87-V93); and α-helix is at α1 (L45-L56). The flexible N- terminal tail (1-21) has been deleted in the representation for the sake of clear visualizations. **b)** Surface representation of Hs-SUMO1 after binding with Hs-Ubc9 enzymes. Salmon color regions in the Hs-SUMO structure represent the residues involved in interaction with Hs-Ubc9. **c,d)** Docked model of Pf-SUMO and E2 enzymes, residues involved in the interaction with Pf-E2 and Hs-E2 enzymes have been highlighted. The Pf-SUMO is shown in blue colour, whereas Pf-E2 and Hs-E2 are shown in green colour. Interacting residues of Pf-SUMO are in salmon colour, and of Pf-E2 and Hs-E2 are in magenta colour.

**Supplementary Fig. 7:**
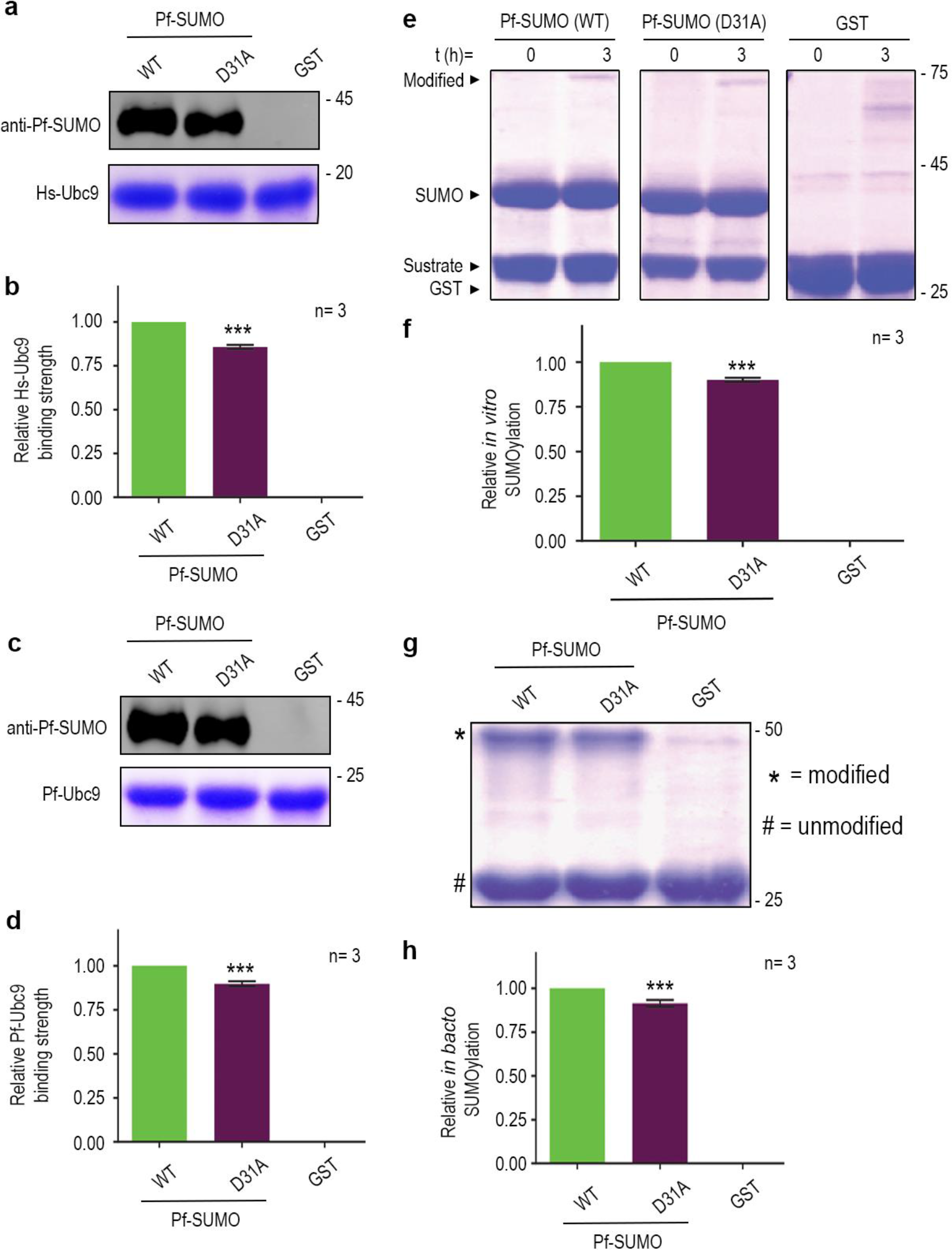
Alanine mutants of Pf-SUMO D31 and D90 mimic wildtype interaction and function patterns. *In vitro* binding assay of Pf-SUMO D31A mutant protein with Hs-Ubc9 **a)**. Upper panels indicate anti-Pf-SUMO antibody blotting, and the coomassie stained lower panels indicate Hs-Ubc9 levels. **b)** The quantification of SUMO binding as seen in (a). **(c and d)** Same as in (a and b); however, the binding is performed over the Pf-Ubc9. GST serves as a negative control. **e)** *In vitro* SUMOylation with Pf-SUMO wild-type and D31A mutant in the presence of purified human SUMOylation machinery components and a standard peptide substrate. **f)** The quantification of *in vitro* SUMOylation as seen in (e). **g)** *In bacto* SUMOylation with Pf-SUMO wild-type and D31A mutant in the presence of human SUMOylation machinery and a standard peptide substrate expressed inside bacteria. **h)** the quantification of the *in bacto* SUMOylation seen in (g). In all reactions, GST serves as a negative control. All statistical analysis was carried out using GraphPad Prism 8.4.3. Column analysis of data sets carried out by One-way ANOVA (nonparametric). Dunnett’s test was used for multiple comparisons. Family-wise significance and confidence level is p<0.001.

**Supplementary Fig. 8:**
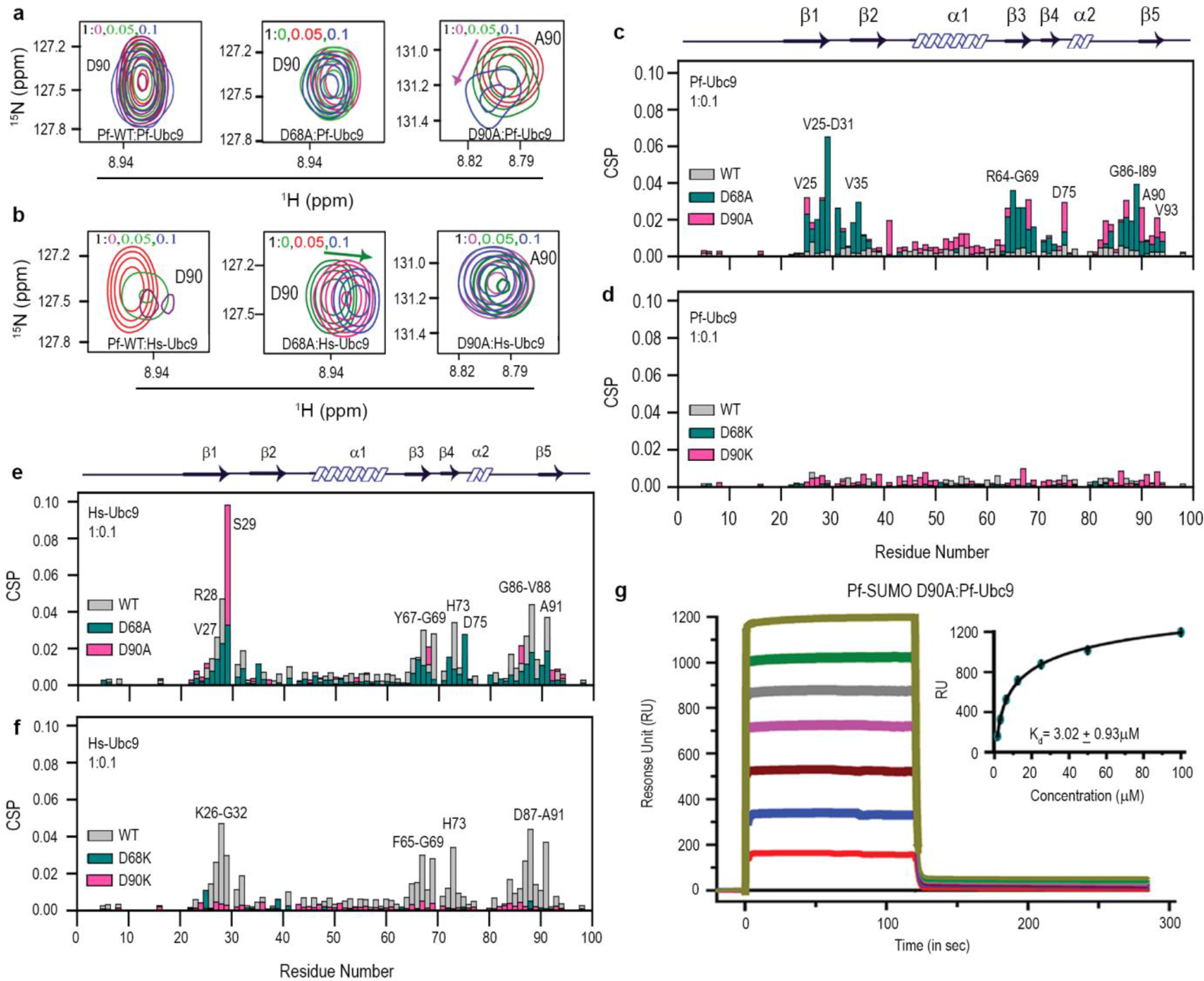
Negatively charged nodes at Aspartate 68 and 90th position govern Pf-SUMO interactions with Ubc9. (**a and b)** The excerpts of selected peaks from Pf-SUMO wildtype, Pf-SUMO D68A, and Pf-SUMO D90A mutants with Pf-Ubc9 and Hs-Ubc9. Arrow indicates the direction of chemical shift perturbations. (**c and d**) Chemical shift perturbations (CSPs) of cross amide peaks obtained from ^15^N-^1^H-HSQC spectra of Pf-SUMO wild type and D68A, D90A and D68K, D90K mutants in the presence of 0.1 equivalent of Pf-Ubc9, respectively. (**e and f**) Same as in (c and d), but Hs-Ubc9 was used instead of Pf-Ubc9. (Gray, *Pf-*SUMO; Pink, D90A & D90K; Dark cyan, D68A & D68K). **g**) Surface plasmon resonance-based observation regarding Pf-SUMO D90A mutant’s interaction with Pf-Ubc9 enzyme. The response unit of Pf-SUMO at different concentrations of Pf-Ubc9 enzyme and curve-fitting by 1:1 binding. The calculated binding affinity (K_d_) is indicated inside the curve.

**Supplementary Fig. 9:**
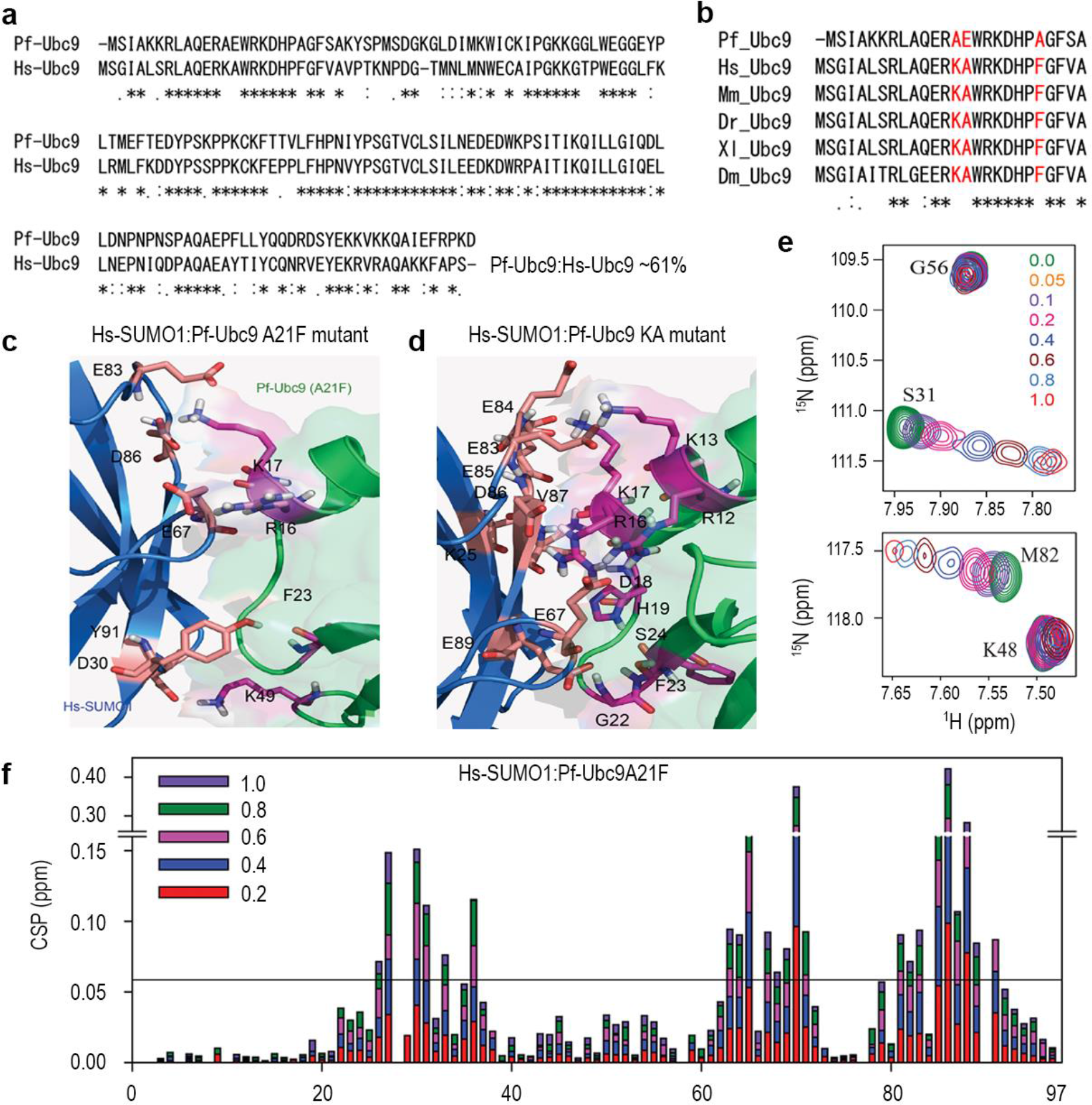
Changes in the key residues in Pf-Ubc9 N-terminus allows interaction with Hs-SUMO1. **a)** Alignment of primary sequences of Plasmodium Ubc9 and human Ubc9 highlighting percentage sequence identity. **b)** Sequence alignment of N-terminal SUMO binding region of Ubc9s from different organisms, including Plasmodium. The key difference in Plasmodium Ubc9 is highlighted in red. Docked model of Hs-SUMO1 and Pf-Ubc9 mutant interaction interface, Pf-Ubc9A21F mutant **c)** and Pf-Ubc9 KA mutant **d)**. The colour coding for Hs-SUMO1 and Pf-Ubc9 mutants are blue and green, respectively. The interacting residues on Hs-SUMO1 are in salmon color, and the Pf-Ubc9 residues are in magenta color. **e)** Overlay ^15^N-^1^H heteronuclear single-quantum coherence (HSQC) spectra of Hs-SUMO1 with Pf-Ubc9 A21F mutant at a ratio of 1: 0.05, 0.1, 0.2, 0.4, 0.6, 0.8, 1.0 i.e. free state to the bound state. Marked and zoomed view of amino acid in HSQC spectra indicates the shifted residues during the interaction. **f)** Quantification of CSPs for residues of Hs-SUMO1 upon titration with Pf-Ubc9 A21F mutant protein.

## Supplementary Tables

**Supplementary Table 1:** Thermodynamics parameters for SUMO-E2 interactions.

| Thermodynamic Parameters | *Pf*-SUMO with *Pf*-E2 | *Pf*-SUMO with *Hs*-E2 | D90A with *Pf*-E2 | *Hs*-SUMO 1 with *Hs*-E2 | *Hs*-SUMO1 with *Pf*-E2 |
| --- | --- | --- | --- | --- | --- |
| Binding Mode | One-sites | One-sites | One-sites | One-sites | One-sites |
| N(no. of sites) | $0.94 \pm 0.006$ | $1.09 \pm 0.036$ | $0.88 \pm 0.025$ | $0.96 \pm 0.003$ | Not quantifiable |
| $K_d$ ($\mu\text{M}$) | $4.03 \pm 0.17$ | $27.42 \pm 1.97$ | $2.50 \pm 0.97$ | $4.18 \pm 0.95$ | Not quantifiable |
| $\Delta H$ (cal/mol) | $-6202 \pm 60.11$ | $-2304 \pm 100.3$ | $-4639 \pm 169.4$ | $-7072 \pm 38.17$ | Not quantifiable |
| $\Delta S$ (cal/mol/deg) | 3.89 | 13.1 | 10.1 | 0.890 | Not quantifiable |

**Supplementary Table 2:** Distance restraints lists and structural statistics of 10 ensemble structures of Pf-SUMO protein.

|  |  |
| --- | --- |
| Distance restraint list |  |
| Sequential | 245 |
| Intra-residual | 367 |
| Medium-range | 342 |
| Long-range | 461 |
| Hydrogen bonds | 40 |
| Dihedral angle restrains ($\phi$ and $\psi$) | 132 |
| Residual restraints violations |  |
| Average no. of distance violations per structure |  |
| 0.1-0.2 Å<sup>0</sup> | 0 |
| 0.2-0.5 Å<sup>0</sup> | 0 |
| > 0.5 Å<sup>0</sup> | 0 |
| Model quality |  |
| RMSD backbone atoms | 0.61 Å<sup>0</sup> |
| RMSD heavy atoms | 1.2 Å<sup>0</sup> |
| RMSD bond lengths | 0.001 Å<sup>0</sup> |
| RMSD bond angles | 0.2 Å<sup>0</sup> |
| Ramachandran plot statistics |  |
| Most favored region (%) | 93.5 |
| Allowed region (%) | 6.5 |
| Additionally allowed region (%) | 0.0 |
| Disallowed (%) | 0.0 |
| Global quality scores (raw/Z score) |  |
| Verify 3D | 0.28/-2.89 |
| Procheck (*phi-psi*) | -0.66/-2.28 |
| Procheck (all) | -0.99/-5.85 |
| MolProbity clash score | 32.32/-4.02 |
| Target function | 9 -10 |
| Z-score | -2.13 |
| Model contents |  |
| Total no. of residues | 98 |
| BMRB accession number | 36011 |
| PDB ID code | 5GJL |

**Supplementary Table 3:**
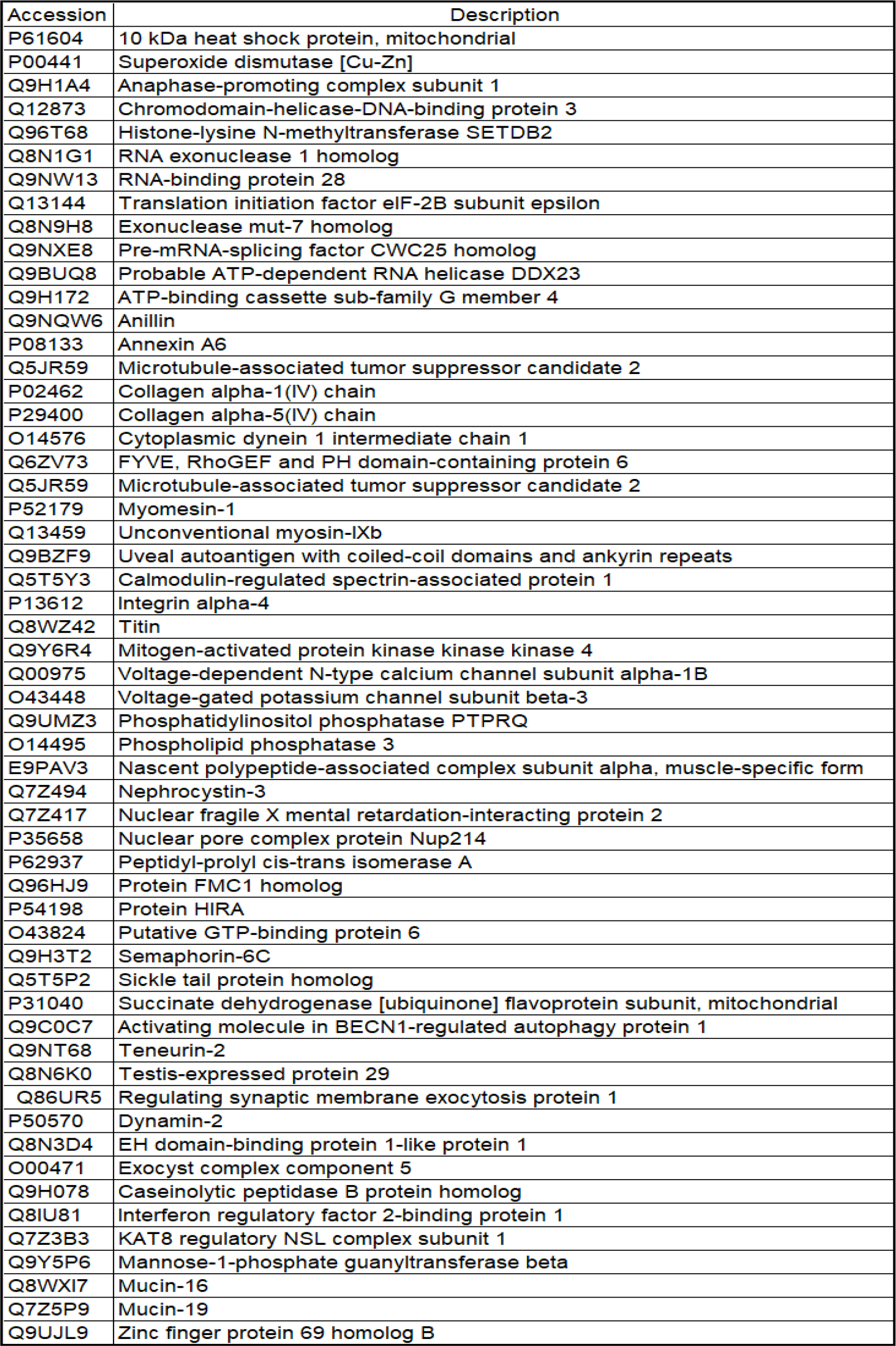
Proteins identified with significant coverage from GFP- trap pull-down using Pf-SUMO and Hs-SUMO1.

